# *Synechococcus* nitrogen gene loss in iron-limited ocean regions

**DOI:** 10.1101/2022.05.24.493279

**Authors:** Garrett Sharpe, Liang Zhao, Meredith G. Meyer, Weida Gong, Shannon M. Burns, Allesandro Tagliabue, Kristen N. Buck, Alyson E. Santoro, Jason R. Graff, Adrian Marchetti, Scott Gifford

## Abstract

*Synechococcus* are the most abundant cyanobacteria in high latitude regions and are responsible for an estimated 17% of annual marine primary productivity. Despite their biogeochemical importance, *Synechococcus* populations have been unevenly sampled across the ocean, with most studies focused on low-latitude strains. In particular, the near absence of *Synechococcus* genomes from high-latitude, High Nutrient Low Chlorophyll (HNLC) regions leaves a gap in our knowledge of picocyanobacterial adaptation to iron limitation and their influence on carbon, nitrogen, and iron cycles. We examined *Synechococcus* populations from the subarctic North Pacific, a well-characterized HNLC region, with quantitative metagenomics. Assembly with short and long reads produced two near complete *Synechococcus* metagenome-assembled genomes (MAGs). Quantitative metagenome-derived abundances of these populations matched well with flow cytometry counts, and the *Synechococcus* MAGs were estimated to comprise >99% of the *Synechococcus* at Station P. Whereas the Station P *Synechococcus* MAGs contained multiple genes for adaptation to iron limitation, both genomes lacked genes for uptake and assimilation of nitrate and nitrite, suggesting a dependence on ammonium, urea, and other forms of recycled nitrogen leading to reduced iron requirements. A global analysis of *Synechococcus* nitrate reductase abundance in the TARA Oceans dataset found nitrate assimilation genes are also lower in other HNLC regions. We propose nitrate and nitrite assimilation gene loss in *Synechococcus* represents an adaptation to severe iron limitation in high-latitude regions where ammonium availability is higher. Our findings have implications for models that quantify the contribution of cyanobacteria to primary production and subsequent carbon export.

**Significance:** The cyanobacterium *Synechococcus* is a major contributor to ocean primary production and biogeochemistry. Here, we used quantitative metagenomics to assemble and enumerate two *Synechococcus* genomes from an iron-limited, High Nutrient Low Chlorophyll region. We show these genomes represent the majority of *Synechococcus* cells at the site and are the first known *Synechococcus* unable to assimilate either nitrate or nitrite. This gene loss is likely due to the high iron quota of these proteins and predominant availability of recycled forms of nitrogen. *Synechococcus’* loss of nitrate assimilation affects their role in elemental cycles (e.g., carbon, nitrogen, and iron), limits their potential for carbon export, and enhances our understanding of *Synechococcus* evolution in response to nutrient limitation and competition.

## Introduction

*Prochlorococcus* and *Synechococcus* are critical components of marine biogeochemical cycles, generating ∼25% of the ocean’s annual net primary production and contributing significantly to carbon export (1-2). *Prochlorococcus* is largely restricted to equatorial and subtropical latitudes, while *Synechococcus* dominates cooler waters in regions of equatorial upwelling and high latitudes (3). Both groups exhibit high levels of strain diversification due to niche specialization arising from variations in environmental conditions (light, temperature, nitrogen, phosphorus, etc.) including iron availability (4-6).

Iron is an essential micronutrient as it is a required cofactor in photosynthetic and respiratory electron transport chains (7, 8). Photosystems I and II require 12 and 3 iron atoms per photosystem, respectively, and the light-harvesting phycobilisome proteins are synthesized by iron-containing enzymes (8). Iron is also required in other key metabolic functions, including nitrate and nitrite assimilation, with nitrate and nitrite reductases requiring 4 and 5 iron atoms per enzyme, respectively (9-11). In High Nutrient Low Chlorophyll (HNLC) regions these cellular iron demands in combination with low iron bioavailability lead to iron limitation of primary production. Three major ocean regions have been identified as HNLC zones: the Equatorial Pacific, the Southern Ocean, and the subarctic North Pacific; together they represent roughly 30% of the world’s oceans (12). These regions are characterized by low phytoplankton biomass and consistently high concentrations of macronutrients in the mixed layer resulting from incomplete utilization due to severe iron limitation (13, 14). Nitrate concentrations for example are in the tens of micromolar range (14).

*Synechococcus* and *Prochlorococcus* strains have evolved diverse adaptations to iron limitation. In non-HNLC regions, *Prochlorococcus* are enriched in iron-storing ferritin genes and iron uptake regulators that enables growth at approximately ten-fold lower iron concentrations and a more rapid response to iron-stress relief (15). In *Synechococcus*, coastal strains possess multiple iron storage, stress regulation, and response genes that are intricately regulated under the dynamic iron conditions of the coastal environment (16). By contrast, pelagic *Synechococcus* strains that grow in the primarily N-limited oligotrophic gyres lack many of these iron-response genes and exhibit a more limited iron regulatory response.

Cyanobacteria, however, are relatively under-sampled in high-latitude HNLC regions, resulting in a major gap in understanding nutrient acquisition and adaptation strategies in these large, biogeochemically important regions. In tropical and subtropical HNLC regions, *Prochlorococcus* and *Synechococcus* have adapted to iron-limiting conditions by substituting iron requiring genes such as Fe-S containing proteins with non-iron containing functional homologues (17, 18, 19). Several *Synechococcus* strains from these regions have high ferritin gene copy numbers for enhanced iron storage (18, 19). Although marker gene analysis has shown that members of the CRD1 clade, a phylogenetically distinct lineage of *Synechococcus* found in the equatorial Pacific, are also present in high-latitude HNLC waters, these regions are not well represented in current bacterial metagenomic datasets, and all currently sequenced isolates, metagenome-assembled genomes (MAGs), and single cell assembled genomes (SAGs) from these clades are derived from low-latitude HNLC zones (19).

The absence of high-latitude HNLC *Synechococcus* genomes leaves a substantial gap in our current knowledge of picocyanobacterial iron adaptation strategies and their importance to biogeochemical cycling. Here, we used quantitative metagenomics and genome assembly to enumerate and characterize cyanobacteria populations at Station P (Ocean Station PAPA) to identify the strategies *Synechococcus* have evolved to succeed in iron-limited HNLC zones and their impact on biogeochemical cycling and carbon export.

## Results

Station P is a low productivity system with high nitrate (7-15 µM) and low iron (< 100 pM) concentrations characteristic of HNLC regions (13, 20, 21). Phytoplankton blooms are rare, and primary production is sustained primarily by intrusion of nutrients from the shallow seasonal pycnocline (22). Fitting with previous observations, phytoplankton at Station P during our sampling consisted primarily of small cells (<5 µm), including *Synechococcus*, small pennate diatoms, and autotrophic flagellates, with low abundance of large (>5 µm) flagellates and diatoms (Fig. 1). Correspondingly, the small size fraction had the highest chlorophyll concentrations, carbon, nitrate, and ammonium uptake rates, and represented 68% of total primary production (Fig. 1). The f-ratios (fraction of total primary production fueled by nitrate: here nitrate uptake / [nitrate + ammonium uptake]) were low for both size fractions, though the small cells had f-ratios half that of the large cells (Fig. 1). Together, the relatively low primary production and f-ratios observed at Station P indicate a system driven by regenerated production, particularly by small phytoplankton cells (23-25).

**Figure 1.**
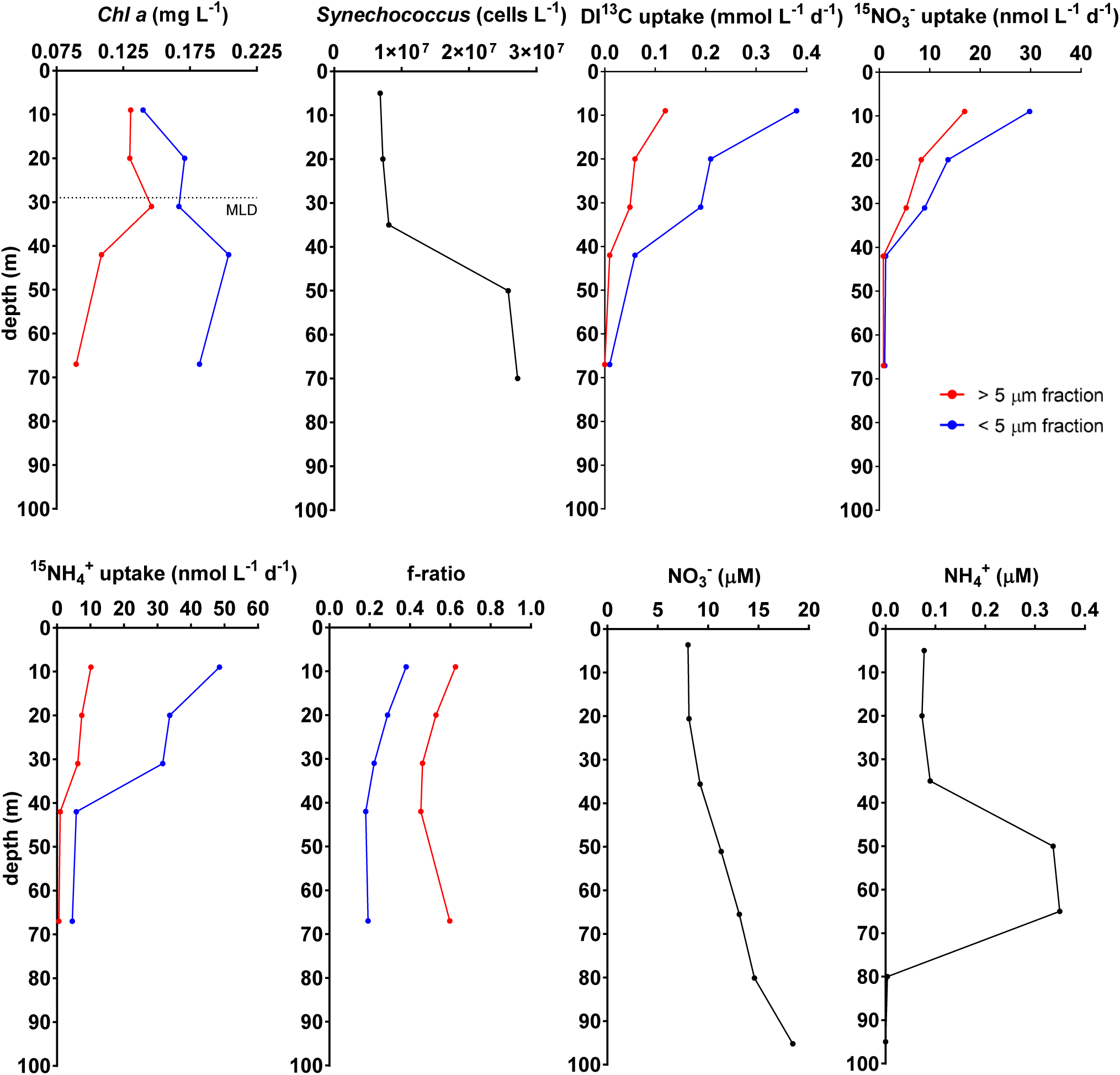
Depth distributions of biological, nutrient, and uptake rates at Station P, September 2018. Chlorophyll a, *Synechococcus* cell concentrations, radiolabeled carbon uptake rates, nitrate uptake rates, ammonia uptake rates. f-ratios calculated from the upper 100 m of the water column for the > 5 µm (red) and < 5 µm (blue) phytoplankton fraction. Non-size-fractionated ammonium NH_4_^+^ and nitrate (NH_4_^+^) concentrations. Average mixed layer depth (MLD) for the cruise (29 m +/-4.5 m) is indicated in *chl-a* graph.

The *Synechococcus-*dominated deep chlorophyll maximum (DCM) was located at 50-70 m, below the mixed layer but well above the ferricline, which was around 200 m. Additionally, iron inputs from dust deposition and mesoscale eddy events are infrequent at Station P compared to the adjacent subtropical North Pacific Gyre, and surface dissolved iron concentrations were low (0.04 ± 0.03 nM during cruise; seasonally ∼0.05 nM spring and summer, ∼0.1 nM winter, 25-29). Ammonium and nitrate concentrations were relatively high at all sampled depths, suggesting neither oxidized nor reduced forms of nitrogen are limiting (Fig. 1 and S1).

We used quantitative metagenomics to enumerate *Synechococcus* abundance at Station P. Genome equivalents were enumerated by identifying single copy recombinase A (*recA*) genes in a metagenome sample, and then converted to volumetric abundances via recovery ratios derived from internal standards added prior to extraction (30-37). A comparison between our quantitative metagenome-derived *Synechococcus* abundances and simultaneously collected flow cytometry *Synechococcus* cell concentrations show strong agreement (Fig. 2A), further supporting the use of internal standard quantitative metagenomics for determining absolute abundances of bacterial groups *in situ. Synechococcus* were the most abundant cyanobacteria at all depths (Fig. 2B), with peak densities of 5 × 10^7^ cells L^−1^ at 50-75 m (Fig. 2B). Taxonomic classification of the *recA* genes revealed the *Synechococcus* community was dominated by two populations, Clade I and IV, both previously found to be abundant at other high-latitude sites (18, 38, 39). These two clades represented >99% of the Station P *Synechococcus* population at all depths, with Clade IV most prevalent at 50 m and Clade I at 70 m.

**Figure 2.**
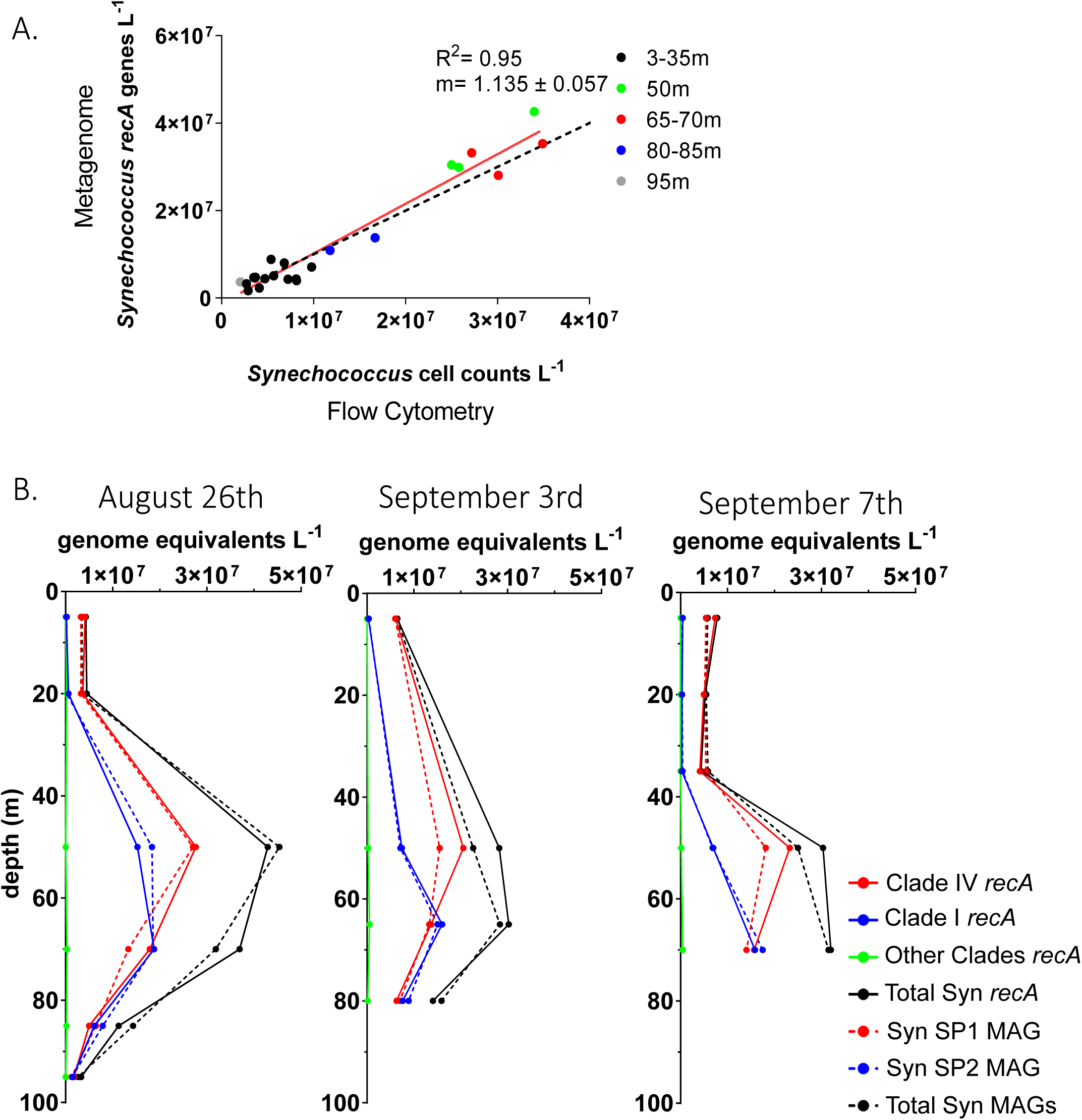
Metagenome-derived volumetric abundances of total *Synechococcus* cells and *Synechococcus* MAG populations at Station P. (A) Comparison of flow cytometry versus metagenome derived *Synechococcus* abundances. The red line is the linear regression model, and the black dashed line is the 1:1 line. (B) Metagenome-derived depth distributions of total *Synechococcus* cells, the subset of *Synechococcus* Clade I and IV populations, and the abundances of cells represented by the three different MAGs. *recA* abundances (solid line) and MAG abundances (dashed line) are separated into total (black), Clade I (blue), Clade IV (red), and Other Clades (green).

Assembly of short and long reads produced two high quality *Synechococcus* genomes (Syn-SP1 and Syn-SP2) representing the two dominant clades (Table 1). Syn-SP1 is most closely related to a Single cell Amplified Genome (SAG; *Synechococcus C sp003208835*) collected from 65 m in the subtropical North Pacific and isolates CC9902 and BL107 from *Synechococcus* Clade IV (40). The second *Synechococcus* MAG (Syn-SP2) is phylogenetically distant from Syn-SP1 (77.3% ANI) and a member of Clade I and is closely related to SAG *Synechococcus*_C sp002500205 and isolates CC9311 and WH8020. Syn-SP1’s genome is similar in size to other known Clade IV members, while Syn-SP2’s genome is smaller than other Clade I members (41). These MAGs are the first genomes from *Synechococcus* clades to dominate these high-latitude regions (18, 39).

**Table 1.** Genome characteristics of the *Synechococcus* Station P (Syn SP) MAGs. Clade: clade of *Synechococcus* MAG binned to within GTDB

| MAG | Source Samples | Clade | Assembled Genome Size (Mbp) | Completeness (%) | Contamination (%) | #Contigs | GC% |
| --- | --- | --- | --- | --- | --- | --- | --- |
| **Syn_SP1** | 0-95m | IV | 1.99 | 95.29 | 0.27 | 116 | 54.87 |
| **Syn_SP2** | 0-95m | I | 2.21 | 97.92 | 1.64 | 81 | 53.4 |

### Absolute quantification of MAG populations

We estimated the volumetric abundances (genomes L^−1^) of the *Synechococcus* MAG populations by deriving a coverage-based recovery ratio from the internal standard genome reads. Surface concentrations of the MAG populations were 3 to 6 × 10^6^ genomes L^−1^, and 2 to 4 × 10^7^ genomes L^−1^ at the DCM (50-70 m) (Fig. 2B). Summed, the two MAGs accounted for nearly all *Synechococcus* genome abundances, as determined by either metagenome-derived *recA* counts (MAGs were 96% of total *Synechococcus recA)* or flow cytometr*y* (97% of *Synechococcus* flow cytometry counts). The Syn-SP1 MAG accounted for an average of 93% of the clade IV population, and the Syn-SP2 MAG accounted for an average of 106% of the clade I population. We further validated the dominance of the SP1 And SP2 populations by mapping the metagenome reads to the MAGS and found 95% of unassembled *Synechococcus* metagenome reads mapped to the two MAGs. The MAGs thus represent the dominant *Synechococcus* populations and their genomic composition at Station P during our sampling.

### Adaptations of *Synechococcus* MAGs

The Station P *Synechococcus* genomes encode several strategies to cope with low iron availability, strategies that are well distributed across *Synechococcus clades* (Fig. 3A and Fig. S2). For iron transport, both genomes possess NRAMP Fe/Mn and *idiABC* iron transporters. In addition, Syn-S1 and Syn-D1 contain an operon for importing iron-chelated siderophores. These siderophore transport genes have previously been identified in a few Clade III, IV, CRD2, and UC-B members (19). It is unclear whether these populations synthesize their own siderophores or can obtain siderophores released by other community members (42). Both Station P genomes possess a single copy of the ferritin gene for iron storage and Fur iron regulatory system. Equatorial HNLC-associated *Synechococcus* clade CRD1 possess multiple ferritin genes, potentially as an adaptation to low iron availability (19). The Station P genomes encode a suite of alternative, low iron-containing proteins for core photosynthesis and electron transport chain functions, in addition to their high-iron dependent counterparts. This includes flavodoxin as a ferredoxin substitute and plastocyanin as a cytochrome c6 substitute, and the presence of only superoxide dismutases that use copper and zinc or nickel cofactors instead of iron (43-45). Overall, the Station P genomes encode multiple strategies for obtaining and conserving iron, but these strategies are not unique to them, rather they are broadly distributed in *Synechococcus* genomes obtained from both low and high iron environments.

**Figure 3.**
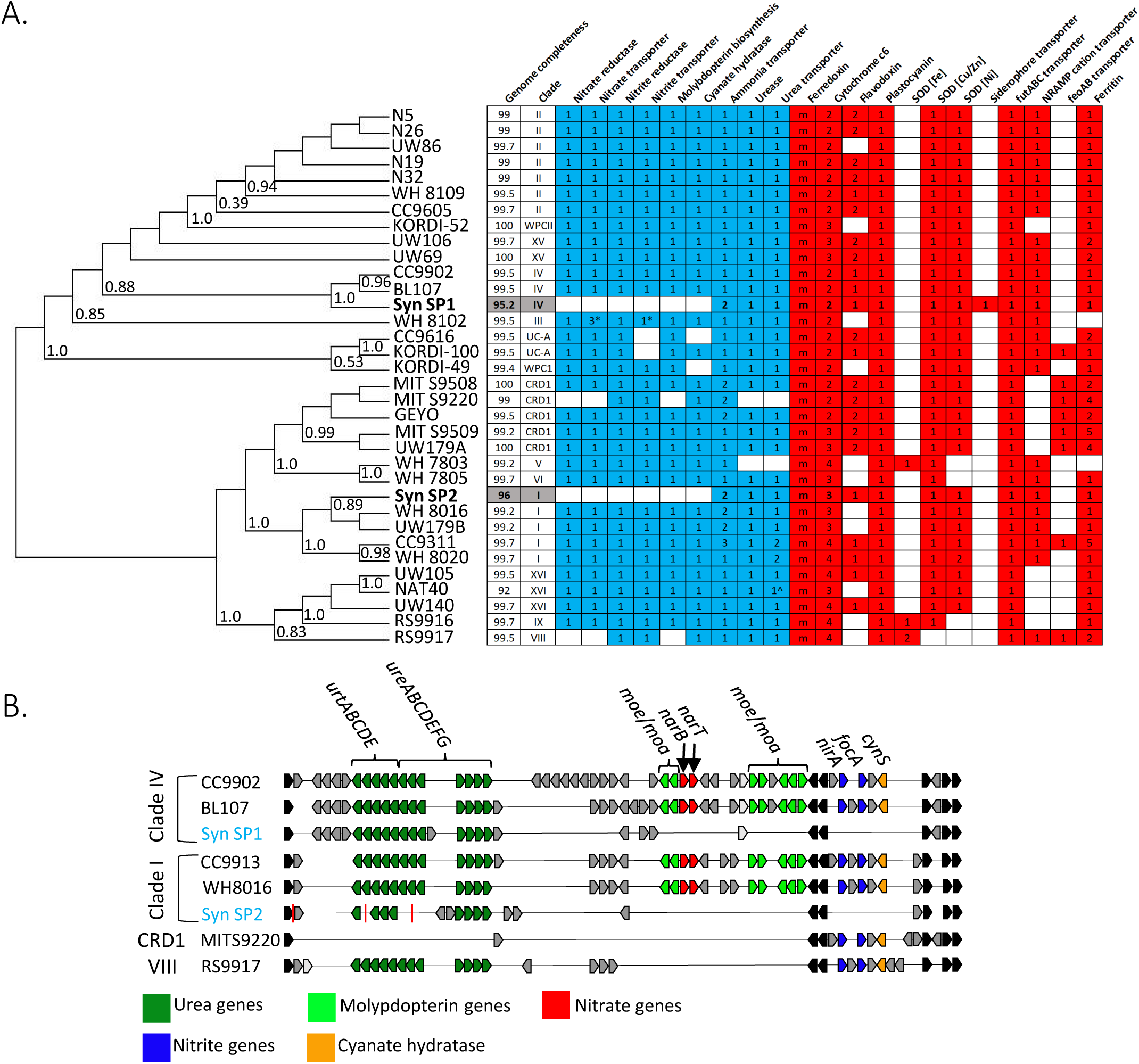
(A) Phylogenomic distribution of nitrogen (blue) and iron (red) associated genes across the *Synechococcus* clade. Only genomes with completeness >90% are included here (for an expanded set of genomes, including partial genomes, see Fig. S3). Phylogenetic relatedness was determined by marker gene comparison using GTDB-Tk. Station P *Synechococcus* MAGs are bolded and highlighted in gray. Numbers in a column represent gene copies in a genome, with ‘m’ standing for multiple copies of the ferredoxin gene, and ‘^’ standing for partial presence of the gene pathway. (B) Distribution of nitrogen assimilation genes across select *Synechococcus* genomes. Station Papa MAGs are labeled in blue. Light gray indicates homologous genes that are present in different order within gene region. Red vertical lines indicate contig breaks.

The Station P *Synechococcus* genomes were unique in their nitrogen assimilation pathway. Within a nitrogen gene cluster highly conserved among cyanobacteria, both genomes are missing genes for nitrate reductase, nitrite reductase, nitrate and nitrite transporters, cyanate hydratase, and the nitrate reductase cofactor molybdopterin biosynthesis genes (Fig. 3B). By contrast, the genomes both contain two distinct ammonium transporters, all urease subunits, and a urea ABC transporter. For the two ammonium transporters found in both Syn-SP1 and Syn-SP2, one is closely related to other *Synechococcus* ammonium transporters, and the other is closely related to euryarchaeal and Thermotogae ammonium transporters (Fig S3).

Support for the absence of nitrate and nitrite utilization genes from Station P *Synechococcus* populations is provided by (1) the high quality and completeness of the MAGs from multiple independent assemblies, (2) the missing nitrogen genes’ location in the interior of a contig flanked by nitrogen genes with homology to taxonomically related genomes, and (3) individual long Nanopore reads lacking these genes (Fig. S4). To further increase confidence that nitrate and nitrite assimilation genes are absent in Station P *Synechococcus* populations, we determined *Synechococcus* nitrogen gene copy numbers in the unassembled metagenomes. If the majority of Station P *Synechococcus* genomes possess nitrate reductase (*narB*), then *narB* copies per genome should be approximately one (Fig. 4). Instead, we found *Synechococcus narB* copy numbers much less than one (range 0.04 to 0.26), supporting their depletion in Station P *Synechococcus* populations. Nitrite reductase (*nirA*) gene ratios were higher (range 0.42 to 1.35), though this may be due to misannotation given *nirA*’s high sequence similarity to sulfite reductase. Copy numbers for the nitrate transporter *narT* and the nitrite/formate transporter *focA* were also much lower than one, supporting depletion in the Station P *Synechococcus* populations. *focA* gene abundance was approximately 25% for the *Synechococcus* community, so it is possible that a *Synechococcus* variant with nitrite reductase represents a fraction of the station P population. By contrast, *Synechococcus* ammonium transporter (*amt*) genome copy numbers were between two and three across all Station P samples (range 0.91 to 2.77), consistent with the two *amt* copies in our *Synechococcus* genomes.

**Figure 4.**
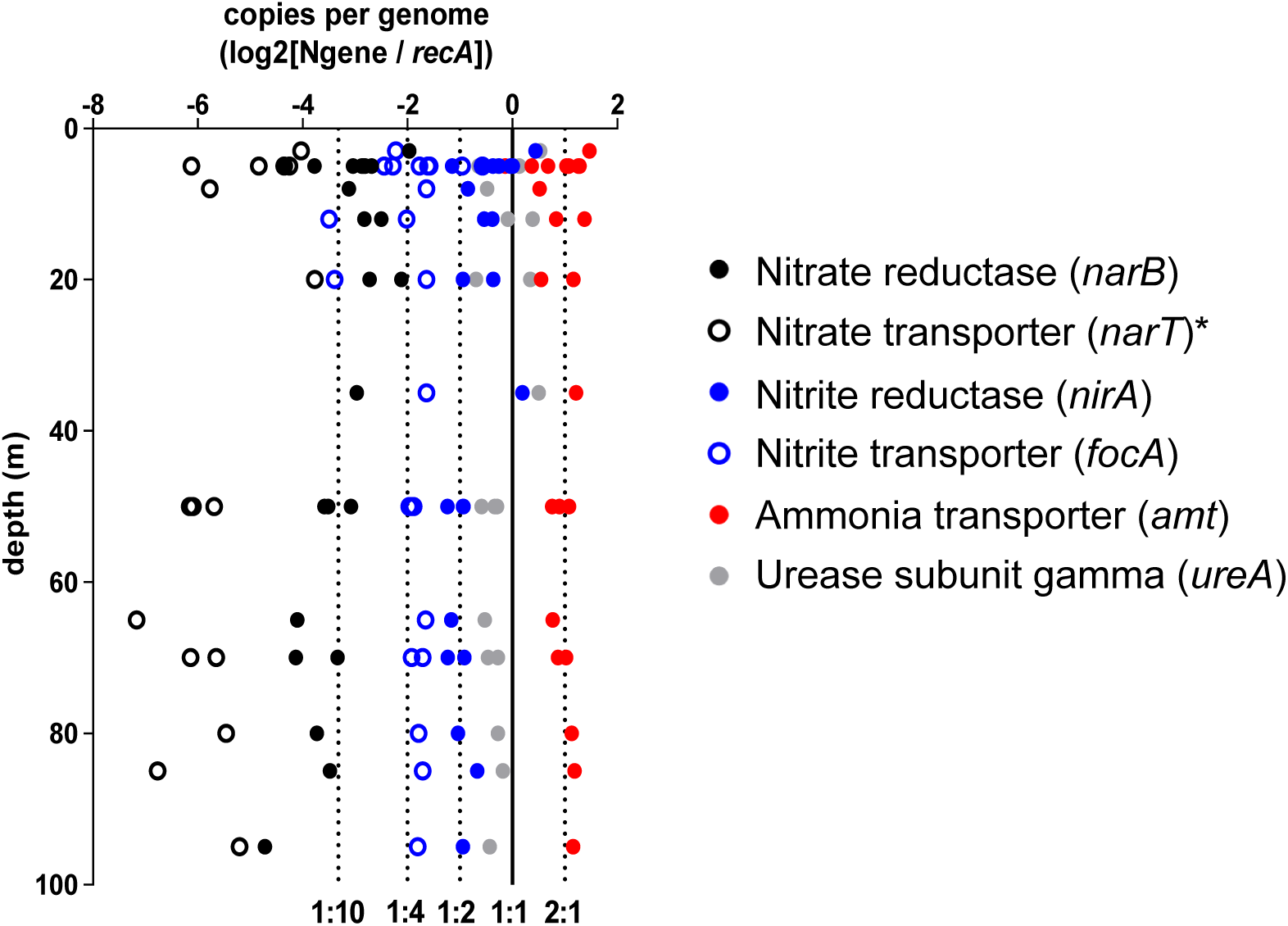
Depth distribution of nitrogen assimilation gene prevalence in Station P unassembled metagenomes. Nitrogen gene copy numbers per genome were calculated as the ratio of *Synechococcus* nitrogen gene read abundance to Synechococcus recA read abundance, and then log2 normalized. The solid vertical line is the 1:1 ratio representing a single copy of the gene per *recA* (*Synechococcus* genome equivalent), with dotted lines representing other genome copy numbers. *Seven samples are not included due to no nitrate transporter reads found in those samples.

The Station P genomes are the first known *Synechococcus* to lack both nitrate and nitrite uptake and utilization genes. Two previously sequenced *Synechococcus* genomes do not encode the ability to use nitrate but are capable of nitrite assimilation: MIT S9508 isolated from an HNLC zone in the Equatorial Pacific, and RS9917 isolated from the Red Sea (46-48). Both isolates were obtained using media with ammonium as the nitrogen source. The loss of both nitrate and nitrite assimilation therefore may be specific to high latitude HNLC *Synechococcus*, though more metagenome sequencing and cultivation on non-traditional *Synechococcus* media (with ammonium used as N source instead of traditionally used nitrate) is needed (49).

The absence of nitrate and nitrite assimilation and the enrichment of ammonium transporters suggests the dominant *Synechococcus* populations at Station P cannot utilize nitrate or nitrite as a nitrogen source and instead rely on ammonium-or other reduced forms of recycled nitrogen for growth. Despite high concentrations of nitrate in this region, the benefit to losing nitrate and nitrite assimilation is a decreased cellular iron and energy demand. Reduction of exogenous nitrate and nitrite to ammonium requires a large quantity of iron, with nitrate reductase containing four iron atoms per enzyme and nitrite reductase containing five iron atoms (11, 50). In many areas of the ocean, iron stress is coupled with nitrogen stress leading cyanobacteria to maintain assimilation capabilities for all sources of N, including nitrate and nitrite, and their associated high iron cost (51). However, in HNLC regions such as Station P, abundant N sources, particularly ammonium and urea, may drive the system more heavily towards iron limitation, resulting in evolutionary pressure to prioritize iron conservation over nitrate utilization via loss of the nitrate/nitrite assimilation pathway (29).

### Global patterns in nitrate and nitrate assimilation loss

To examine whether *Synechococcus* nitrate and nitrite assimilation loss is widespread in the global ocean, we extended the nitrogen gene copy number analysis to the TARA Oceans metagenomes. TARA Oceans contigs from the to 1.6 µm size fraction were annotated via a BLASTX search to identify contigs containing *Synechococcus* nitrate reductase (*narB*), nitrite reductase (*nirR*), ammonium transporter (*amt*), and recombinase A (*recA*) genes. These contigs were paired with their corresponding TARA-generated gene coverage data to calculate nitrogen assimilation and utilization gene copy numbers per genome equivalents. The ratios at each station were then compared to surface nitrate and dissolved iron concentrations (N:Fe ratios) predicted by the PISCES global ocean biogeochemical model (52, 53). Station P *Synechococcus narB* genome copies were depleted compared to most TARA stations (Fig. 5A). Nitrite reductase (*nirR*) genome copy numbers were not significantly different between TARA stations, though Station P samples from the DCM or lower had *nirR* copy numbers below most TARA sites. By contrast, *Synechococcus* ammonium transporters (*amtT*) were typically found at greater than one copy per genome at both Station P and TARA stations, though there was a clear enrichment at Station P (Fig 5A). Based on the TARA metagenome and PISCES nutrient data, nitrate reductase (*narB)* copy numbers are low in the HNLC regions of the southeastern and equatorial Pacific that have high nitrate and low chlorophyll *a* concentrations (Fig. 5B and 5C). The lowest *narB* copy numbers correspond with Station P and the TARA Stations with the highest N:Fe ratios, all of which are within or adjacent to HNLC regions, supporting nitrate assimilation gene loss is linked to iron limitation (Fig 5C).

**Figure 5.**
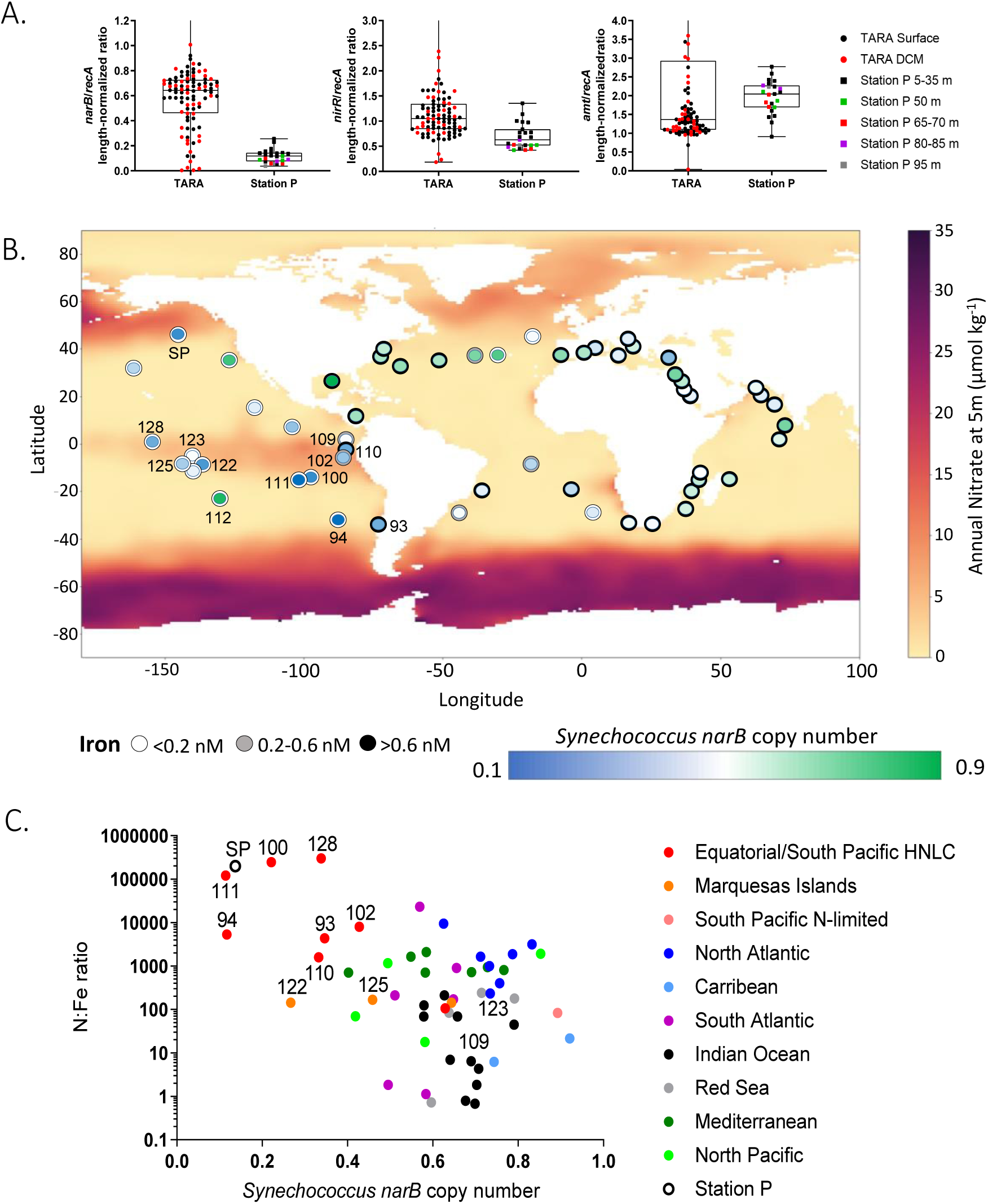
(A) Gene copy numbers for nitrate reductase, nitrite reductase, and ammonia transporters at TARA Ocean and Station P sites. (B) Global patterns of nitrate (background map) and dissolved iron concentrations (outer circle of each datapoint), and *Synechococcus* nitrate reductase genome copy numbers (inner circle of each datapoint) for TARA stations and Station P (large circle). *Synechococcus* nitrogen copy numbers were calculated using gene length normalized ratios derived from the publicly available TARA dataset. Annual nitrate concentrations were derived from the WOA2018 global dataset. Dissolved iron concentrations determined from the PISCES biogeochemical model and BYONIC 3R biogeochemistry dataset. (C) Nutrient ratios versus *Synechococcus* nitrate reductase gene copy number of the TARA stations and EXPORTS Station P. Select stations within the equatorial Pacific HNLC zone and the Eastern Tropical Southern Pacific are labeled with their TARA ID.

## Discussion

Our findings show the dominant *Synechococcus* at Station P are incapable of assimilating either nitrate or nitrite and instead rely on reduced nitrogen sources such as ammonium and urea. The loss of genes for both nitrate and nitrite assimilation reduce the cellular iron demand required by their respective reductases, imparting a fitness advantage in this low-iron HNLC region. The f-ratios were low at Station P, particularly in small cells below the mixed layer where *Synechococcus* is most abundant, which can be attributed to the lack of *Synechococcus* nitrate and nitrite uptake capabilities. In other regions, *Synechococcus* populations have lost nitrate assimilation while retaining nitrite assimilation. The TARA Oceans nitrogen gene analysis indicates *Synechococcus* nitrate reductases are also depleted within the equatorial and sub-tropical Southern Pacific HNLC regions. The results suggest having both nitrate and nitrite assimilation capabilities or nitrite assimilation alone is not a core trait of marine *Synechococcus*.

The disparate loss of nitrogen assimilation genes across the *Synechococcus* phylogeny suggests multiple factors might be influencing their retention. If iron limitation exerts a significant selective pressure for nitrogen gene loss, why have high latitude HNLC *Synechococcus* lost both nitrate and nitrite assimilation while equatorial HNLC *Synechococcus* strains only abandon a portion of the pathway? This may be due to dueling pressures between iron stress and competition for reduced nitrogen species. Complete loss of nitrate and nitrite assimilation may only be possible at high latitudes where there is reduced competition for ammonium from other cyanobacteria. *Prochlorococus* is a major competitor for reduced nitrogen species, often relying solely on ammonium as their nitrogen source. They are dominant in many low to mid latitude regions but absent at high latitudes (3, 6, 11, 54), including Station P where they are undetectable by pigment analysis, flow cytometry, and metagenomics. In low latitude HNLC regions, competition with *Prochlorococus* and heterotrophic bacteria may result in *Synechococcus* at least maintaining nitrite uptake. In high latitude HNLC regions, competition for ammonium is lower due to the absence of *Prochlorococcus*. Further, overall community competition for ammonium is likely low at Station P given the relatively high standing stocks of ammonium at the DCM; Station P’s peak ammonium concentration was greater than threefold higher than those measured in equatorial HNLC sites (Fig. S5).

The model for *Prochlorococcus* nitrogen gene gain and loss appears to be a close counterpart to *Synechococcus*, where nitrogen gene retention is selected for by nitrogen competition but selected against in iron-or light-limiting environments (11, 55). Low-light *Prochlorococcus* typically possess only nitrite assimilation genes and are abundant at the subsurface nitrite maximum at the base of the euphotic zone (54). Competition for ammonium near the nitrite maximum is intense due to the presence of ammonia-oxidizing archaea, driving low-light *Prochlorococcus* to acquire nitrite generated by ammonia oxidizers (56-58). Additionally, most low-light *Prochlorococcus* are restricted to the base of the euphotic zone, where the energy required to reduce nitrate may drive selection against maintaining nitrate assimilation (11, 54). High-light *Prochlorococcus* suffer less light limitation and many strains possess both nitrate and nitrite assimilation pathways (6, 11, 59). Critically, high-light *Prochlorococcus* clades adapted to low-iron conditions in the Equatorial Pacific HNLC zone do not contain nitrate and nitrite utilization genes, reducing their iron requirements (11, 17, 60).

The inability to assimilate nitrate and nitrite by *Synechococcus* populations suggests a re-evaluation of their role in HNLC nutrient cycles and carbon export. In these regions, iron availability limits primary production and the degree to which nitrate can be utilized due to the iron requirements for nitrate reduction and assimilation. New iron delivered to this region via lateral advection, entrainment from subsurface waters, or from atmospheric inputs leads to pulses of nitrate-based new production typically dominated by large phytoplankton (i.e., diatoms) that exceed surface remineralization and can be exported (61). However, according to our findings, any new inputs of iron would not stimulate nitrate utilization and subsequent growth in the dominant *Synechococcus* population at Station P, prohibiting them from significantly contributing to organic carbon export fluxes. Instead, Station P *Synechococcus* are confined to a tight recycling loop where they are dependent on other primary producers to assimilate nitrate, which then directly or indirectly regulates the subsequent availability of recycled forms of nitrogen such as ammonium or urea. The direct inability of *Synechococcus* to bloom when pulses of iron are added to the system means their fixed carbon is continuously cycled in the surface ocean when grazed upon or remineralized by bacterioplankton, leaving little carbon to be exported to depth. Thus, whereas small cyanobacteria such as *Prochlorococcus* and *Synechococcus* can be major contributors to ocean carbon export, in the North Pacific and possibly other HNLC regions, this does not appear to be the case (1, 6, 62, 63).

While our analysis suggests the dominant *Synechococcus* populations at Station P do not contain nitrate and nitrite assimilation pathways, we did detect the presence of these genes at relatively low levels (< 10% of genome equivalents) in the unassembled metagenome. This suggests there may be a small population of nitrate utilizing *Synechococcus* maintained in the community. At different times of the year these populations may increase in abundance, potentially when iron is pulsed into the system altering nitrogen cycles and carbon export potential. This emphasizes the need for further genomic and biogeochemical investigations to model the role of cyanobacteria more accurately in the Fe, N, and C cycles of HNLC zones. Further, if nitrogen assimilation adaptation driven by iron limitation is prevalent within other HNLC *Synechococcus* populations, as our TARA Oceans analysis suggests, it has significant impacts on nutrient cycling and associated carbon export by the cyanobacterial community in a sizable fraction of the global ocean. Overall, our results further support how iron availability affects primary productivity directly through limitation but also has fundamentally shaped phytoplankton functional capabilities leading to cascading effects on marine biogeochemical cycles and food webs.

## Materials and Methods

Samples were collected at Station P in the North Pacific in August and September 2018 as part of the NASA Export Processes in the Ocean from RemoTe Sensing (EXPORTS) expedition (24). Flow cytometric enumeration of *Synechococcus* was performed on a Becton Dickinson Influx Cell Sorter (BD-ICS) flow cytometer while at sea following previously published protocols (64). For uptake incubations, seawater was collected with a trace metal clean rosette, aliquoted into acid washed bottles, and inoculated with NaH^13^CO_3_ isotope and Na^15^NO_3-_ isotope at 10% of ambient DIC and NO_3-_concentrations and incubated for 24 hours. Samples were gravity filtered through pre-combusted GF/F filters, dried and stored until onshore analysis at the UC Davis Stable Isotope Facility (23). Ambient nutrient concentrations were collected as described in Siegel et al. (2021; ref 24).

For the metagenomes, seawater was prefiltered through a 5 µm membrane filter and cells collected on 0.2 µm membrane filter. DNA for short read sequencing was extracted from the filters using a DNeasy Powerwater kit. Internal genomic standards were added immediately before starting the extraction. Metagenomes were sequenced with HiSeq 4000 as 2 × 150 bp reads. Metagenome reads were annotated with a DIAMOND search against the NCBI Refseq protein database. Metagenome assembled genomes (MAGS) were assembled with metaspades and then mapped for read coverage with Bowtie2. Contigs were each binned by MetaBAT, MaxBin, and CONCOCT, and then consolidated using DAS Tool and CheckM. *Synechococcus* MAG quality was increased by performing several additional assemblies including Oxford nanopore long reads and manual curation. DNA for long read sequencing was obtained by phenol chloroform extraction.

*Synechococcus* volumetric abundances were derived from internal standard normalized metagenomes by recA recovery and internal standard genome recovery. Metagenome recA genes were identified by a DIAMOND homology search against a custom RecA protein database, and the resulting *recA* read counts were converted to volumetric abundances using the internal standards recovery ratio and volume of seawater filtered. To calculate MAG population abundances, a coverage-based ratio was calculated for each of the three internal standard genomes by mapping metagenome reads to the internal standard reference genomes. The mean depth of coverage was calculated by dividing the total number of bases mapped by the size of the genomes. This mean depth of coverage represents the number of internal standard genomes recovered through sequencing and was divided by the number of genome molecules added to the sample to determine the recovery ratio. The *Synechococcus* MAG volumetric abundances was then determined by mapping metagenome reads onto the *Synechococcus* MAG, calculating the mean depth of coverage, and then dividing by the coverage-based recovery ratio and volume of seawater filtered. Detailed descriptions of sample collection, processing, and analysis can be found in the supplemental methods.

## Acknowledgements

(NASA), chief scientist Deborah Steinberg (VIMS), the captain and crew of the *R/V Roger Revelle*, and all the members of the EXPORTs team. We thank Matt Kellom for Amt alignments and analysis. This research was supported by NASA grant 80NSSC17K0552 to S.G., A.M, and Nicolas Cassar, NASA award 80NSSC18K1431 to A.S., NASA award 80NSSC17K0568 to Michael J. Behrenfeld (Co-I J.R. Graff), and NSF award OCE-1756433 to K.N.B.

## References

1. T. L. Richardson, G. A. Jackson, Small phytoplankton and carbon export from the surface ocean. Science 315, 838–840 (2007).

2. P. Flombaum, et al., Present and future global distributions of the marine Cyanobacteria Prochlorococcus and Synechococcus. Proc. Natl. Acad. Sci. U. S. A. 110, 9824–9829 (2013).

3. Z. I. Johnson, et al., Niche partitioning among Prochlorococcus ecotypes along ocean-scale environmental gradients. Science 311, 1737–1740 (2006).

4. J. A. Sohm, D. G. Capone, Phosphorus dynamics of the tropical and subtropical north Atlantic: Trichodesmium spp. versus bulk plankton. Mar. Ecol. Prog. Ser. 317, 21–28 (2006).

5. N. A. Ahlgren, G. Rocap, Diversity and Distribution of Marine Synechococcus: Multiple Gene Phylogenies for Consensus Classification and Development of qPCR Assays for Sensitive Measurement of Clades in the Ocean. Front. Microbiol. 3, 213 (2012).

6. A. C. Martiny, S. Kathuria, P. M. Berube, Widespread metabolic potential for nitrite and nitrate assimilation among Prochlorococcus ecotypes. Proc. Natl. Acad. Sci. U. S. A. 106, 10787–10792 (2009).

7. J. A. Raven, M. C. W. Evans, R. E. Korb, The role of trace metals in photosynthetic electron transport in O2-evolving organisms. Photosynth. Res. 60, 111–150 (1999).

8. J. Morrissey, C. Bowler, Iron utilization in marine cyanobacteria and eukaryotic algae. Front. Microbiol. 3, 43 (2012).

9. J. A. Raven, The iron and molybdenum use efficiencies of plant growth with different energy, carbon and nitrogen sources. New Phytol. 109, 279–287 (1988).

10. F. M. M. Morel, N. M. Price, The biogeochemical cycles of trace metals in the oceans. Science 300, 944–947 (2003).

11. P. M. Berube, A. Rasmussen, R. Braakman, R. Stepanauskas, S. W. Chisholm, Emergence of trait variability through the lens of nitrogen assimilation in Prochlorococcus. Elife 8 (2019).

12. P. W. Boyd, M. J. Ellwood, The biogeochemical cycle of iron in the ocean. Nat. Geosci. 3, 675–682 (2010).

13. J. H. Martin, R. M. Gordon, S. Fitzwater, W. W. Broenkow, Vertex: phytoplankton/iron studies in the Gulf of Alaska. Deep Sea Res. A 36, 649–680 (1989).

14. J. K. Moore, S. C. Doney, D. M. Glover, I. Y. Fung, Iron cycling and nutrient-limitation patterns in surface waters of the World Ocean. Deep Sea Res. Part 2 Top. Stud. Oceanogr. 49, 463–507 (2001).

15. A. W. Thompson, K. Huang, M. A. Saito, S. W. Chisholm, Transcriptome response of high-and low-light-adapted Prochlorococcus strains to changing iron availability. ISME J. 5, 1580–1594 (2011).

16. K. R. M. Mackey, et al., Divergent responses of Atlantic coastal and oceanic Synechococcus to iron limitation. Proc. Natl. Acad. Sci. U. S. A. 112, 9944–9949 (2015).

17. D. B. Rusch, A. C. Martiny, C. L. Dupont, A. L. Halpern, J. C. Venter, Characterization of Prochlorococcus clades from iron-depleted oceanic regions. Proc. Natl. Acad. Sci. U. S. A. 107, 16184–16189 (2010).

18. J. A. Sohm, et al., Co-occurring Synechococcus ecotypes occupy four major oceanic regimes defined by temperature, macronutrients and iron. ISME J. 10, 333–345 (2016).

19. N. A. Ahlgren, B. S. Belisle, M. D. Lee, Genomic mosaicism underlies the adaptation of marine Synechococcus ecotypes to distinct oceanic iron niches. Environ. Microbiol. 22, 1801–1815 (2020).

20. F. A. Whitney, Nutrient variability in the mixed layer of the subarctic Pacific Ocean, 1987– 2010. J. Oceanogr. 67, 481–492 (2011).

21. J. Nishioka, H. Obata, Dissolved iron distribution in the western and central subarctic Pacific: HNLC water formation and biogeochemical processes. Limnol. Oceanogr. 62, 2004–2022 (2017).

22. P. Boyd, P. J. Harrison, Phytoplankton dynamics in the NE subarctic Pacific. Deep Sea Res. Part 2 Top. Stud. Oceanogr. 46, 2405–2432 (1999).

23. M. G. Meyer, W. Gong, S. Kafrissen, O. Torano, D. Varela, A. E. Santoro, N. Cassar, S. M. Gifford, A. K. Niebergall, G. C. Sharpe, A. Marchetti, in revision, Phytoplankton size-class contributions to new and regenerated production during the EXPORTS Northeast Pacific Ocean field deployment. Elem Sci Anth (2022).

24. D. A. Siegel, et al., An operational overview of the EXport Processes in the Ocean from RemoTe Sensing (EXPORTS) Northeast Pacific field deployment. Elem Sci Anth 9, 00107 (2021).

25. H. McNair, M. Meyer, S. Lerch, A. E. Maas, B. Stephens, J. Fox, K. N. Buck, S. M. Burns, Cetinic, M. Cohn, C. Durkin, S. Gifford, W. Gong, J. R. Graff, B. Jenkins, E. L. Jones, A. E. Santoro, C. H. Shea, K. Stamieszkin, D. K. Steinberg, A. Marchetti, C. A. Carlson, S. Menden-Deuer, M. A. Brzezinski, D. A. Siegel, T. Rynearson, submitted, A quantitative analysis of food web dynamics in the Subarctic Pacific reveals a regenerative system with low export potential. Elementa: Science of the Anthropocene (2022).

26. R. C. Hamme, et al., Volcanic ash fuels anomalous plankton bloom in subarctic northeast Pacific. Geophys. Res. Lett. 37 (2010).

27. P. Xiu, A. P. Palacz, F. Chai, E. G. Roy, M. L. Wells, Iron flux induced by Haida eddies in the Gulf of Alaska. Geophys. Res. Lett. 38 (2011).

28. J. N. Fitzsimmons, et al., Daily to decadal variability of size-fractionated iron and iron-binding ligands at the Hawaii Ocean Time-series Station ALOHA. Geochim. Cosmochim. Acta 171, 303–324 (2015).

29. P. J. Harrison, Station Papa Time Series: Insights into Ecosystem Dynamics. J. Oceanogr. 58, 259–264 (2002).

30. S. M. Gifford, S. Sharma, J. M. Rinta-Kanto, M. A. Moran, Quantitative analysis of a deeply sequenced marine microbial metatranscriptome. ISME J. 5, 461–472 (2011).

31. S. M. Gifford, J. W. Becker, O. A. Sosa, D. J. Repeta, E. F. DeLong, Quantitative Transcriptomics Reveals the Growth- and Nutrient-Dependent Response of a Streamlined Marine Methylotroph to Methanol and Naturally Occurring Dissolved Organic Matter. MBio 7 (2016).

32. S. M. Gifford, et al., Microbial Niche Diversification in the Galápagos Archipelago and Its Response to El Niño. Front. Microbiol. 11, 575194 (2020).

33. B. M. Satinsky, S. M. Gifford, B. C. Crump, M. A. Moran, Use of internal standards for quantitative metatranscriptome and metagenome analysis. Methods Enzymol. 531, 237– 250 (2013).

34. B. M. Satinsky, et al., Microspatial gene expression patterns in the Amazon River Plume. Proc. Natl. Acad. Sci. U. S. A. 111, 11085–11090 (2014).

35. B. M. Satinsky, et al., Expression patterns of elemental cycling genes in the Amazon River Plume. ISME J. 11, 1852–1864 (2017).

36. Y. Lin, S. Gifford, H. Ducklow, O. Schofield, N. Cassar, Towards Quantitative Microbiome Community Profiling Using Internal Standards. Appl. Environ. Microbiol. 85 (2019).

37. E. Crossette, et al., Metagenomic Quantification of Genes with Internal Standards. Mbio 12 (2021).

38. K. Zwirglmaier, et al., Global phylogeography of marine Synechococcus and Prochlorococcus reveals a distinct partitioning of lineages among oceanic biomes. Environ. Microbiol. 10, 147–161 (2008).

39. M. L. Paulsen, et al., Synechococcus in the Atlantic Gateway to the Arctic Ocean. Frontiers in Marine Science 3 (2016).

40. P. M. Berube, et al., Single cell genomes of Prochlorococcus, Synechococcus, and sympatric microbes from diverse marine environments. Sci Data 5, 180154 (2018).

41. M. D. Lee, et al., Marine Synechococcus isolates representing globally abundant genomic lineages demonstrate a unique evolutionary path of genome reduction without a decrease in GC content. Environ. Microbiol. 21, 1677–1686 (2019).

42. J. J. Morris, R. E. Lenski, E. R. Zinser, The Black Queen Hypothesis: evolution of dependencies through adaptive gene loss. MBio 3 (2012).

43. D. L. Erdner, N. M. Price, G. J. Doucette, M. L. Peleato, D. M. Anderson, Characterization of ferredoxin and flavodoxin as markers of iron limitation in marine phytoplankton. Mar. Ecol. Prog. Ser. 184, 43–53 (1999).

44. C. Frazão, et al., Ab initio determination of the crystal structure of cytochrome c6 and comparison with plastocyanin. Structure 3, 1159–1169 (1995).

45. Y. Sheng, et al., Superoxide dismutases and superoxide reductases. Chem. Rev. 114, 3854–3918 (2014).

46. L. R. Moore, A. F. Post, G. Rocap, S. W. Chisholm, Utilization of different nitrogen sources by the marine cyanobacteria Prochlorococcus and Synechococcus. Limnol. Oceanogr. 47, 989–996 (2002).

47. B. S. Belisle, et al., Genome Sequences of Synechococcus sp. Strain MIT S9220 and Cocultured Cyanophage SynMITS9220M01. Microbiol Resour Announc 9 (2020).

48. N. J. Fuller, et al., Clade-specific 16S ribosomal DNA oligonucleotides reveal the predominance of a single marine Synechococcus clade throughout a stratified water column in the Red Sea. Appl. Environ. Microbiol. 69, 2430–2443 (2003).

49. J. B. Waterbury, The cyanobacteria—isolation, purification and identification. The prokaryotes 4, 1053–1073 (2006).

50. M. M. Dose, M. Hirasawa, S. Kleis-SanFrancisco, E. L. Lew, D. B. Knaff, The ferredoxin-binding site of ferredoxin: Nitrite oxidoreductase. Differential chemical modification of the free enzyme and its complex with ferredoxin. Plant Physiol. 114, 1047–1053 (1997).

51. M. A. Saito, et al., Multiple nutrient stresses at intersecting Pacific Ocean biomes detected by protein biomarkers. Science 345, 1173–1177 (2014).

52. C. Richon, A. Tagliabue, Biogeochemical feedbacks associated with the response of micronutrient recycling by zooplankton to climate change. Glob. Chang. Biol. 27, 4758– 4770 (2021).

53. Y. Shaked, B. S. Twining, A. Tagliabue, M. T. Maldonado, Probing the bioavailability of dissolved iron to marine eukaryotic phytoplankton using in situ single cell iron quotas. Global Biogeochem. Cycles 35 (2021).

54. P. M. Berube, et al., Physiology and evolution of nitrate acquisition in Prochlorococcus. ISME J. 9, 1195–1207 (2015).

55. L. J. Ustick, et al., Metagenomic analysis reveals global-scale patterns of ocean nutrient limitation. Science 372, 287–291 (2021).

56. P. M. Berube, A. Coe, S. E. Roggensack, S. W. Chisholm, Temporal dynamics of Prochlorococcus cells with the potential for nitrate assimilation in the subtropical Atlantic and Pacific oceans. Limnol. Oceanogr. 61, 482–495 (2016).

57. M. W. Lomas, F. Lipschultz, Forming the primary nitrite maximum: Nitrifiers or phytoplankton? Limnol. Oceanogr. 51, 2453–2467 (2006).

58. W. Martens-Habbena, et al., The production of nitric oxide by marine ammonia-oxidizing archaea and inhibition of archaeal ammonia oxidation by a nitric oxide scavenger. Environ. Microbiol. 17, 2261–2274 (2015).

59. R. R. Malmstrom, et al., Temporal dynamics of Prochlorococcus ecotypes in the Atlantic and Pacific oceans. ISME J. 4, 1252–1264 (2010).

60. R. R. Malmstrom, et al., Ecology of uncultured Prochlorococcus clades revealed through single-cell genomics and biogeographic analysis. ISME J. 7, 184–198 (2013).

61. R. T. Pollard, et al., Southern Ocean deep-water carbon export enhanced by natural iron fertilization. Nature 457, 577–580 (2009).

62. J. R. Casey, M. W. Lomas, J. Mandecki, D. E. Walker, Prochlorococcus contributes to new production in the Sargasso Sea deep chlorophyll maximum. Geophys. Res. Lett. 34 (2007).

63. L. Guidi, et al., Plankton networks driving carbon export in the oligotrophic ocean. Nature 532, 465–470 (2016).

64. J.R. Graff, M. J. Behrenfeld, Photoacclimaon Responses in Subarcc Atlanc Phytoplankton Following a Natural Mixing-Restraficaon Event. Frontiers in Marine Science 5 (2018).

