## Supplemental Information for "*Synechococcus* nitrogen gene loss in iron-limited ocean regions"

### SUPPLEMENTAL MATERIALS and METHODS

**Sample Collection and Preservation.** Samples were collected at Station P in the North Pacific from August 18th to September 7th, 2018 as part of the NASA Export Processes in the Ocean from RemoTe Sensing (EXPORTS) expedition (1). Discrete water samples from 5 to 95 m were collected using a CTD/rosette and dispensed into acid-washed 4 L bottles. For each sample, 1 to 2.5 L of water was peristaltically pumped through a 25 mm diameter 5.0 µm pore TMTP Isopore filter and a 25mm diameter 0.2 µm pore PES. Filters were placed into a 2 mL cryovial using sterilized forceps and flash-frozen in liquid nitrogen.

**Flow Cytometric Determinations of *Synechococcus*.** Analysis of seawater samples for phytoplankton communities were performed on a Becton Dickinson Influx Cell Sorter (BD-ICS) flow cytometer while at sea following previously published protocols (2). Briefly, the ICS used a blue laser (488 nm) and two detectors for fluorescence (692 ± 40 nm and 530 ± 40 nm) and two for scattering properties (forward scatter and side scatter). The ICS optical alignment was performed daily with fluorescent beads (Spherotech, SPHEROTM ~3.0 µm Ultra Rainbow Calibration Particles) following standard protocols and photomultiplier tube (PMT) gains were identical for all samples to minimize artifacts due to instrument settings. Samples for flow cytometric analysis were collected from CTD rosette mounted Niskin bottles and analyzed within ~30 min of collection. At Station P, an average of >5000 phytoplankton cells were interrogated per sample and specific groups were identified based on light scattering and fluorescence properties. *Synechococcus* have a distinct scattering and fluorescence signature that allow these autotrophic cyanobacteria to easily be identified in relation to other microalgal groups. Cell concentrations are calculated from cell counts determined over a known amount of time (cells per second) and a sample flow rate (volume per second) determined for each set of samples collected from each CTD cast.

**Isotopic Uptake Incubations.** Triplicate samples for carbon and nitrogen stable isotope incubations were collected via a trace metal clean CTD rosette in 1 liter acid-cleaned polycarbonate bottles. Each bottle was spiked with a dual inoculation of $NaH^{13}CO_3$ isotope and $Na^{15}NO_3^-$ isotope at a concentration of approximately 10% of ambient concentrations of DIC and $NO_3^-$, respectively. Bottles were incubated for 24 hours. Following incubation, two of the triplicate samples were gravity filtered through a 5 µm polycarbonate filter with the filtrate subsequently being vacuum filtered through a pre-combusted Whatman glass fiber filter (GF/F) to collect size fractionated samples. Sample material retained on the polycarbonate filter was washed onto a new, pre-combusted GF/F using filtered (0.2 µm) seawater. One sample was retained as a whole sample and filtered through a pre-combusted GF/F. Filters were dried and stored until onshore preparation and analysis at the UC Davis Stable Isotope Facility. Ambient nutrient concentrations used in this analysis were collected according to Siegel et al. (2021). For further details on carbon and nitrogen stable isotopic uptake methods refer to Meyer et al., *in revision*.

**DNA Extraction and Internal Standard Addition.** DNA was extracted from the filters using a DNeasy Powerwater kit (Qiagen, Hilden, Germany). The 0.2 µm filters were removed from the storage tubes using sterilized forceps and placed in Powerwater bead tube. PW1 lysis buffer was added to the original filter storage tube, vortexed briefly, and then transferred to the Powerwater bead tube to ensure complete transfer of sample. For quantitative analysis, three bacterial genomic internal standards (*Blautia producta*, *Deinococcus radiodurans*, and *Thermus thermophilus*), were individually added at 4 ng each (previously quantified by PicoGreen) in 25 µl volumes to the Powerwater bead tube prior to starting the DNA extraction, aiming for 1% of total DNA being comprised of genomic standards and assuming 1000 ng native DNA in the sample. The bead tube was shaken with a vortex adapter for 5 minutes. The remainder of the extraction followed the kit manual, except for an additional ethanol removal step where the filter column was transferred to a clean 2mL microcentrifuge tube and spun at 13,000 rcf for 2 minutes to remove trace ethanol prior to eluting the DNA. The DNA was eluted in 100 µl of elution buffer (10 mM Tris, pH 8.0). DNA yields were quantified with the Quant-iT PicoGreen dsDNA kit (Molecular Probes Inc., Organ, United States).

**Library Preparation and Sequencing.** Metagenomic libraries were prepared using the KAPA HyperPlus Library Preparation Kit (Kapa Biosystems Scientific, Massachusetts, United States). 100 ng of purified DNA diluted in 10 mM Tris-HCL was used as input. For the fragmentation step, a 23 minute fragmentation time at 37 °C produced an average DNA fragment size of 330 bp (including ligated adapters). Samples were barcoded via the addition of a NimbleGen SeqCap Adapters (Roche NimbleGen Inc., Wisconsin, United States; containing single index barcodes) at a concentration of 1 µM. Reaction cleanup steps were conducted using AMPure XP reagent beads (Beckman Coulter, California, United States) with a 0.8X SPRI bead cleanup following adapter ligation and a 1X cleanup following library amplification. Additionally, a dual-SPRI bead size selection selecting for 400 bp fragments was performed after the post-ligation cleanup. Five cycles were used for the library amplification reaction. Library fragment composition and quality was analyzed on the Agilent 2100 Bioanalyzer with High Sensitivity DNA chip (Agilent Technologies, Waldbronn, Germany) and quantified with the Quant-iT PicoGreen dsDNA kit (Molecular Probes Inc., Organ, United States). Equimolar concentrations of the barcoded libraries were then pooled and sequenced using HiSeq 4000 platform 150 bp PE (Illumina, San Diego, CA).

**DNA extraction for long read sequencing.** A phenol chloroform extraction was performed on one sample (65 m, September 5$^{th}$) to generate long DNA fragments for Oxford Nanopore sequencing. Cells on the filter were resuspended in 500 µl of $T_{50}E_{50}$ (50 mM Tris pH 7.5, 50 mM EDTA pH 8) and frozen at -80°C overnight with the filter remaining in the tube. Lysis was performed via the addition of 25 µL lysozyme (10 mg lysozyme per 1mL 10mM Tris, pH 7.5) to the tube with the frozen pellet and thawed in a room temperature water bath. The cell lysate was frozen again at -80°C overnight and thawed the following day in a room temperature water bath. Protein was degraded by adding 100 µl STE (0.5% SDS in $T_{50}E_{50}$) and 35 µl of proteinase K (2 mg proteinase K per 1 mL DNase-free water) and incubating for an hour at 55˚C with occasional mixing. RNA was removed via the addition of 5 µl RNase A (10 mg RNase A per 1 mL DNase-free water) and incubating for 30 min at 37°C, with mixing by inversion done at 10-minute intervals. Deproteinization was carried out through the addition of 200 µl of 5 M sodium perchlorate. 640 µl of Tris-equilibrated phenol:chloroform (1:1 mixture of phenol and chloroform with phenol buffered to pH 7.8-8.0) was added to the sample, mixed by inversion and separated into the organic and aqueous phases via centrifugation at 12,000 rpm for 10 minutes. The aqueous phase was transferred to a new tube, and the phenol:chloroform separation was repeated. The aqueous phase was once again transferred to a new tube, and 600 µl chloroform was mixed into the aqueous phase, followed by centrifugation at 12,000 rpm for 10 minutes. The aqueous phase was transferred to a new tube, and the chloroform addition and centrifugation was repeated. To concentrate the extracted DNA, the aqueous phase was transferred to a new tube, and a 1/10$^{th}$ volume of 3 M sodium acetate was mixed in with the aqueous layer. Two volumes of cold 100% ethanol were mixed with the sample, and the reaction placed on ice for 30 minutes. The precipitated DNA was pelleted by centrifugation at 12,000 rpm for 10 minutes. The supernatant was removed by pipette. One milliliter of 70% ethanol was added to the pellet, spun briefly, and removed by pipette. The pellet was air-dried and then resuspended in 50 µL of molecular-grade water. DNA concentration was quantified as 5.5 ng $\mu L^{-1}$ with the Quant-iT PicoGreen dsDNA kit (Molecular Probes Inc., Organ, United States), and the presence of larger fragments (~10kb) was determined with the Agilent High Sensitivity DNA kit (Agilent Technologies, Waldbronn, Germany). Forty microliters of this DNA sample was sent to the UNC High-throughput Sequencing Facility for sequencing on an Oxford Nanopore flow cell.

**Metagenome Read Processing**. For Illumina sequences, FastQC was performed to determine the general quality of the raw reads (3). Trimmomatic (paired-end mode) was used to remove adaptor sequences and low-quality base pairs (bp) from both the forward and reverse reads with a sliding window looked of 10 base pairs and trimming base pairs that had an average PHRED score of less than 20 (4). Trimmed reads < 50 bp were removed. Another FastQC was performed to determine the quality of the trimmed reads and ensure that adaptor sequences were successfully removed. Paired forward and reverse trimmed reads were merged using Pear (--p-value 0.01 --min-overlap 10 --min-asm-length 50 --min-trim-length 50 --quality-threshold 0 --max-uncalled-base 1.0 --test-method 1 --empirical-freqs --score-method 2 --cap 40) a minimum overlap of 10 bp to be successfully merged into one assembled read (5). The merged reads were quality checked with FastQC. The merged reads, unmerged forward reads, and unpaired forward and reverse reads that lost their paired read during trimming were all concatenated into one file to be used for read annotations. The concatenated file metagenome reads were then annotated with a DIAMOND search against the NCBI Refseq protein database (version 95; default settings) (6).

As described above, three genomic internal standards were added at a known concentration to each sample prior to DNA extraction. Each internal standard represents a genome alien to surface ocean communities. Internal standard reads were identified via a BLASTn search (megablast; e-value <0.001, >90% identity) of the processed metagenome reads against the internal standard genomes, and results filtered to retain hits that had a >95% identity to internal standard sequences. To identify the number of internal standard gene hits, a BLASTx of the BLASTn-identified reads was used. All confirmed internal standard reads were removed from the dataset before proceeding with analysis.

For the Nanopore reads, *NanoPlot* was used to determine the quantity, size, and quality of reads (7). Sequencing adapters were removed from the reads with *PoreChop*. *Nanofilt* was used to trim the edges of each read to ensure removal of adaptors (--headcrop 50, --tailcrop 50), and removed reads with an average quality score <8 and/or a length of < 100 bp (-l 100, -q 8) (6). Quality, quantity, and read length of the quality-controlled and Porechopped reads were assessed using *NanoPlot* (7).

**MAG assembly and annotation.** metaSPAdes was used to assemble contigs from surface reads (all samples from 5-35 m) and deep reads (50-95 m) separately (8). Metagenome reads were mapped onto contigs with Bowtie2 (9), and the resulting alignments were indexed and sorted using SAMtools (10). Metagenome binning was then performed using MetaBAT, MaxBin, and CONCOCT (11-13). Bins from the three binning programs were consolidated using DAS Tool, and the quality and general characteristics of each bin were assessed with CheckM (14, 15). Taxonomic assignments were given to the MAGs (or bins) based on Average Nucleotide Identity (ANI) and placement on the reference phylogenomic tree using GTDB-Tk (version 0.3.2; Genome Taxonomy Database release 04-RS89) (16).

To improve the quality of the *Synechococcus* MAGs, several additional assemblies were performed: 1) nanopore deep and surface samples reads, 2) *Synechococcus*-annotated Illumina reads with (all?) nanopore reads, and 3) Illumina reads from individual samples. For MAG Syn_SP1, we used minimus2 to further assemble the bin reconstructed from the *Synechococcus*-annotated Illumina with nanopore assembly (min identity 0.94; min overlap 40bp) (17). For Syn_SP2, we ran minimus2 on the bins from the *Synechococcus* Illumina with nanopore assembly and the deep assembly. We specifically focused on the nitrogen assimilation gene cluster region. However, SP2's N gene region was still broken into 3 contigs despite the fact that nanopore long reads spanned across them and suggested there were 2 ~400bp gaps. We then manually pulled contigs from the deep samples with nanopore assembly and the FN446 assembly (single sample) to fill in those gaps. Contigs that did not get assembled by minimus2 were mapped onto the assembled contigs with minimap2 and visualized with IGV (18,19) to identify and remove redundant contigs. The resulting bins were visualized with Anvi'o (v5) to inspect contigs' coverage and composition (20). After manual curation, the MAGs were assessed by checkM to see if quality had improved.

For each bin, protein-coding genes were identified and annotated with Prokka (gene calls made using Prodigal) (21). Blastp searches were performed against the Prokka protein sequence files using query sequences related to nitrogen and iron metabolism to identify the functional capabilities of the *Synechococcus* MAGs.

***recA* quantification and taxonomy**. *Synechococcus* genome equivalents for each sample were estimated with absolute *recA* gene count. The *recA* gene encodes a DNA repair protein recombinase A and is a well-described single-copy gene (ref). To identify recA genes in the metagenome samples, a protein database containing viral sequences and sequences annotated with the key words "recombinase RecA," "protein RecA," "recombinase A," or "RecA protein" was assembled from the NCBI RefSeq protein database (v.95). Metagenome reads were then compared to this custom database using a DIAMOND homology search. To reduce the chance of getting false positives, top hits with a bit score >50 were counted as a recA gene only if the read was also annotated as a recA gene in the homology search against the NCBI RefSeq database. To directly compare recA-based abundances with MAG abundances, the recA reads were re-annotated with a GTDB RecA database containing RecA sequences from the GTDB representative genomes and the *Synechococcus* MAG RecA sequences (22).

The resulting *recA* read counts for each *Synechococcus* clade were converted to volumetric *recA* abundances using equations described in (23, 24):

$$1)\ S_r = \frac{S_S}{S_P}$$

$$2)\ R_r = \frac{S_r}{S_a}$$

$$3)\ G_a = \frac{G_S}{R_r}$$

(1) $S_r$: Copies of internal standard genome recovered in sequence library.
 $S_S$: protein encoding internal standard reads in the sequence library.
 $S_p$: protein encoding genes in the internal standard reference genome.
(2) $R_r$: read-based recovery ratio. The proportion of standard molecules added that were sequenced.
 $S_a$: molecules of internal standard genome added to the sample.
(3) $G_a$: Molecules in the sample of any gene category.
 $G_s$: total reads of any gene category in the sample sequence library

For each internal standard genome, the number of gene hits was divided by the total number of genes in the genome to get a value for the number of internal standard genomes recovered. To convert metagenome reads to absolute values, a conversion factor for a given sample was created by dividing the number of internal standard genomes added by the number of internal standard genomes recovered. The average conversion factor for a sample was derived from the average of the resulting three conversion factor values. recA reads recovered in the sample was then multiplied by this conversion factor and normalized by dividing by the volume filtered to get the value for absolute number of recA reads per liter, which is a proxy for cell abundance.

**Whole genome quantification.** *Synechococcus* MAG volumetric genome abundances were estimated with a coverage-based recovery ratio derived from the internal standard genomes with the following calculations:

$$4)\ R_{cov} = \frac{S_{cov}}{S_a}$$

$$5)\ M_a = \frac{M_{cov}}{R_{cov}}$$

(4) $R_{cov}$: coverage-based recovery ratio.

$S_{cov}$: mean depth of coverage of internal standard genomes by metagenomic reads.

(5) $M_a$: molecules of any genome (MAG) in the sample.

$M_{cov}$: mean depth of coverage of any genome (MAG) in the sequence library.

Metagenomic reads were mapped onto the internal standard genomes with bowtie2 and the mean depth of coverage was calculated by dividing the total number of bases mapped by the size of the genomes (SAMtools bedcov) (9, 10). The mean depth of coverage represents the number of internal standard genome recovered through sequencing, and this was divided by the number of genomes added to the samples to get at the recovery ratio. The number of MAGs recovered from the sequence library was retrieved by mapping reads onto the MAG and calculating mean depth of coverage. The volumetric abundance of the MAGs was determined by normalizing MAG abundance in the sample (Ma) by the volume of seawater filtered.

**Genome and gene alignment analysis.** Contigs for the clade IV and clade I MAGs were reordered based on the *Synechococcus* reference genomes BL107 and CC9311 respectively using Mauve (25). Annotations and gene locations for the reference genomes and the MAGs were produced as a gbk file by Prokka, and the annotated reordered MAGs were compared with the annotated reference genomes to identify whether the gaps for the missing nitrogen genes were located within or at the edge of the MAG contigs. GBK files of only the region in which the N genes are present for most *Synechococcus* species were generated by Prokka for the MAGs and the reference genomes BL107 (IV), CC9902 (IV), WH8016 (I), CC9311 (I), MIT S9220 (CRD1), and RS9917 (VIII). The nitrogen gene regions were then visualized and compared with Clinker (26). Nanopore long reads were mapped onto the MAGs with minimap2 and coverage of the nitrogen gene regions was visualized with IGV (18,19).

**Nitrate reductase gene abundance and global data comparison.** *Synechococcus* nitrate reductases in the EXPORTS samples were identified and quantified using keyword searches ('*Synechococcus*' and 'nitrate reductase') within the diamond results of each EXPORTS sample. *Synechococcus* nitrate reductases were grouped by *Synechococcus* clade in order to compare their abundance to the *Synechococcus* MAG abundance and *Synechococcus recA* abundance.

For comparison of *Synechococcus* nitrate reductase gene frequencies between our sample and TARA samples, the NR to *recA* ratio (nitrate reductases per *Synechococcus* genome equivalent) was used. The 0.2 µm fraction gene contig and gene profile datasets from the TARA database was used to generate NR to *recA* ratios for other ocean regions to determine whether other areas exhibit similar nitrate reductase abundance profiles to those of Station PAPA. A diamond blast was performed on the TARA station contig dataset OM-RGC_v2 (nucleic acid converted to amino acid database), and *Synechococcus* nitrate reductase and *recA* sequences were selected using keyword searches. The gene coverages of these selected genes were extracted from the OM-RGC-v2_gene_profile_metaG TARA file, and used to calculate NR: *recA*. Nitrate reductase associated proteins were not included when calculating the NR to *recA* ratio.

Iron and nitrate concentrations for each TARA station were calculated using the PISCES biogeochemistry model in order to relate NR: *recA* to nitrate and iron availability at each station (27, 28).

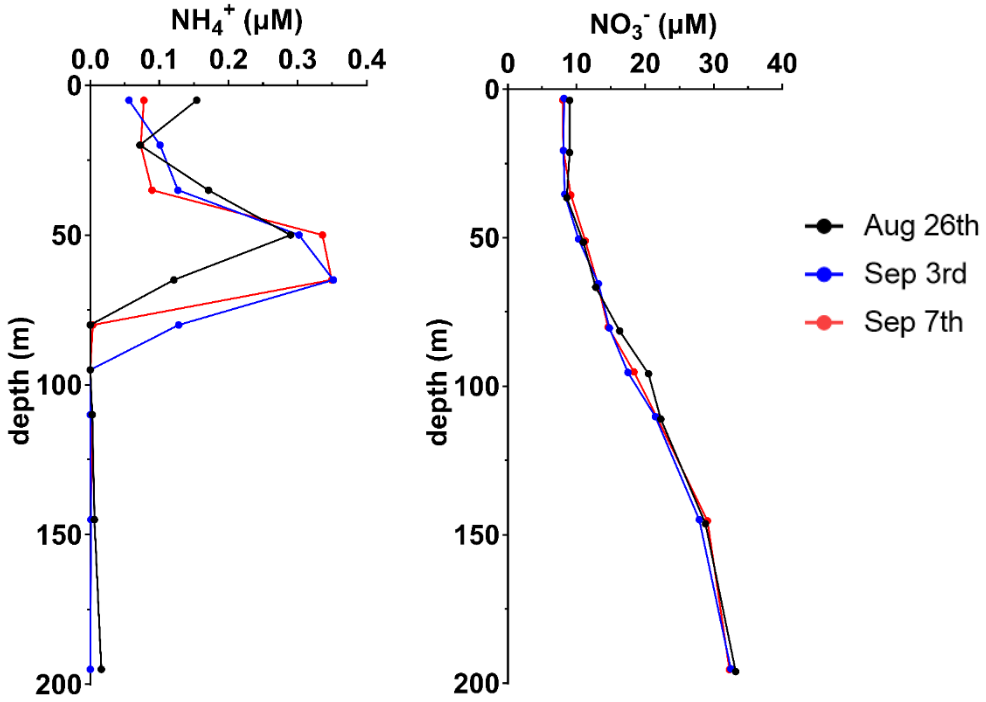


Figure S1. Ammonium and nitrate concentrations (μM) from 0-200 m in depth on August 26th (black), September 3rd (blue), and September 7th (red).

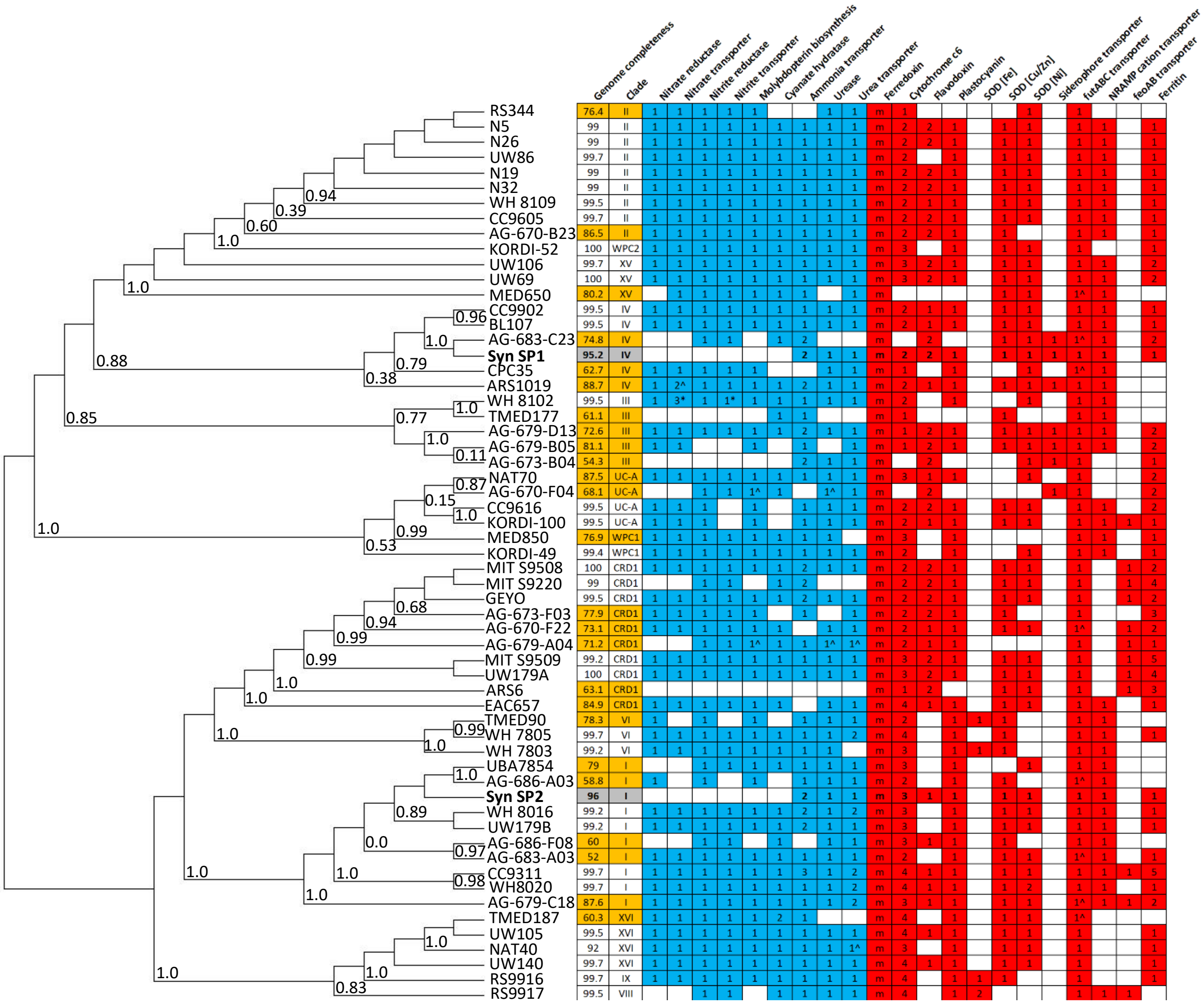


Figure S2. Phylogenomic distribution of nitrogen (blue) and iron (red) associated genes across all analyzed genomes of the *Synechococcus* clade. Station PAPA *Synechococcus* MAGs are bolded and highlighted in gray. Numbers in a column represent gene copies in a genome, with m standing for multiple copies of the ferredoxin gene. Genomes with an estimate completeness of <90% are included here and highlighted in orange. * represent the inclusion of an active Nitrate/nitrite ABC transporter for WH 8102. ^ represent the presence of a partially complete pathway, where only some genes for that function were present within genomes with low completeness.

*A.*

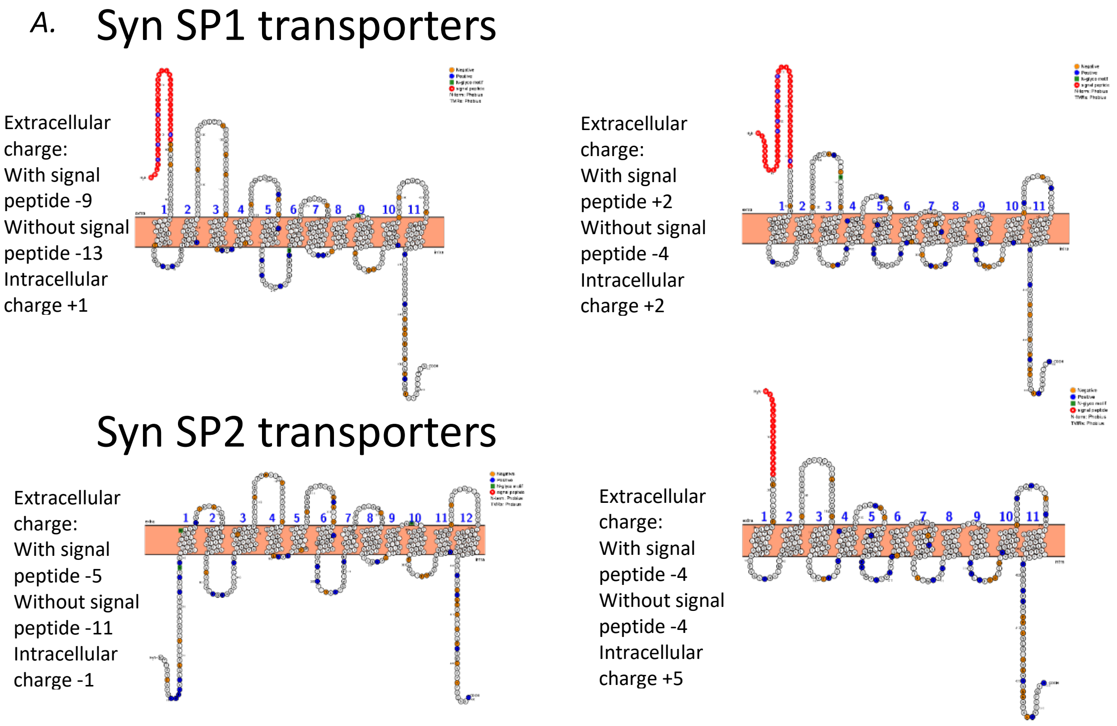


*B.*

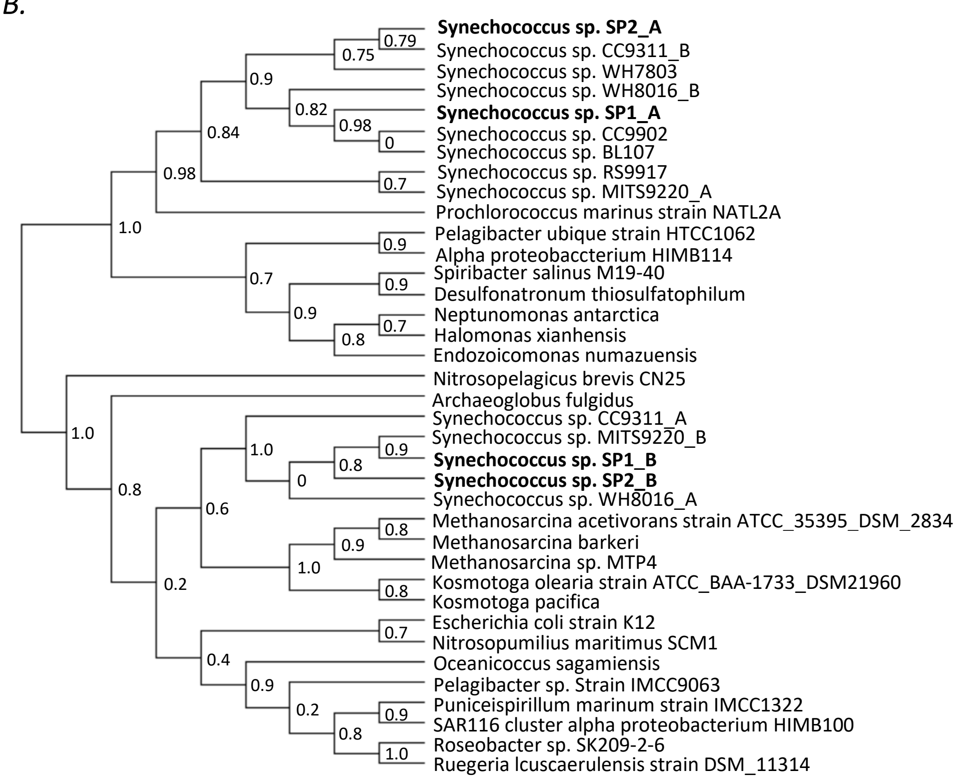


Figure S3. A. Structure of Syn SP1 and SP2 ammonium transporters predicted using Protter (29). Syn SP2 ammonium transporter A likely has a long signal motif that got mistaken as a 12th transmembrane region. B. Phylogenetic tree of ammonium transporters. Multiple sequence alignments of ammonia transporter amino acid sequences were performed with MUSCLE, and GBlocks removed poorly aligned and divergent regions of the sequence alignments and identified 174 positions to compare for phylogenetic analysis. PhyML generated the phylogenetic tree from the sequence alignments using an Approximate Likelihood-Ratio Test (Alrt) for branch support.

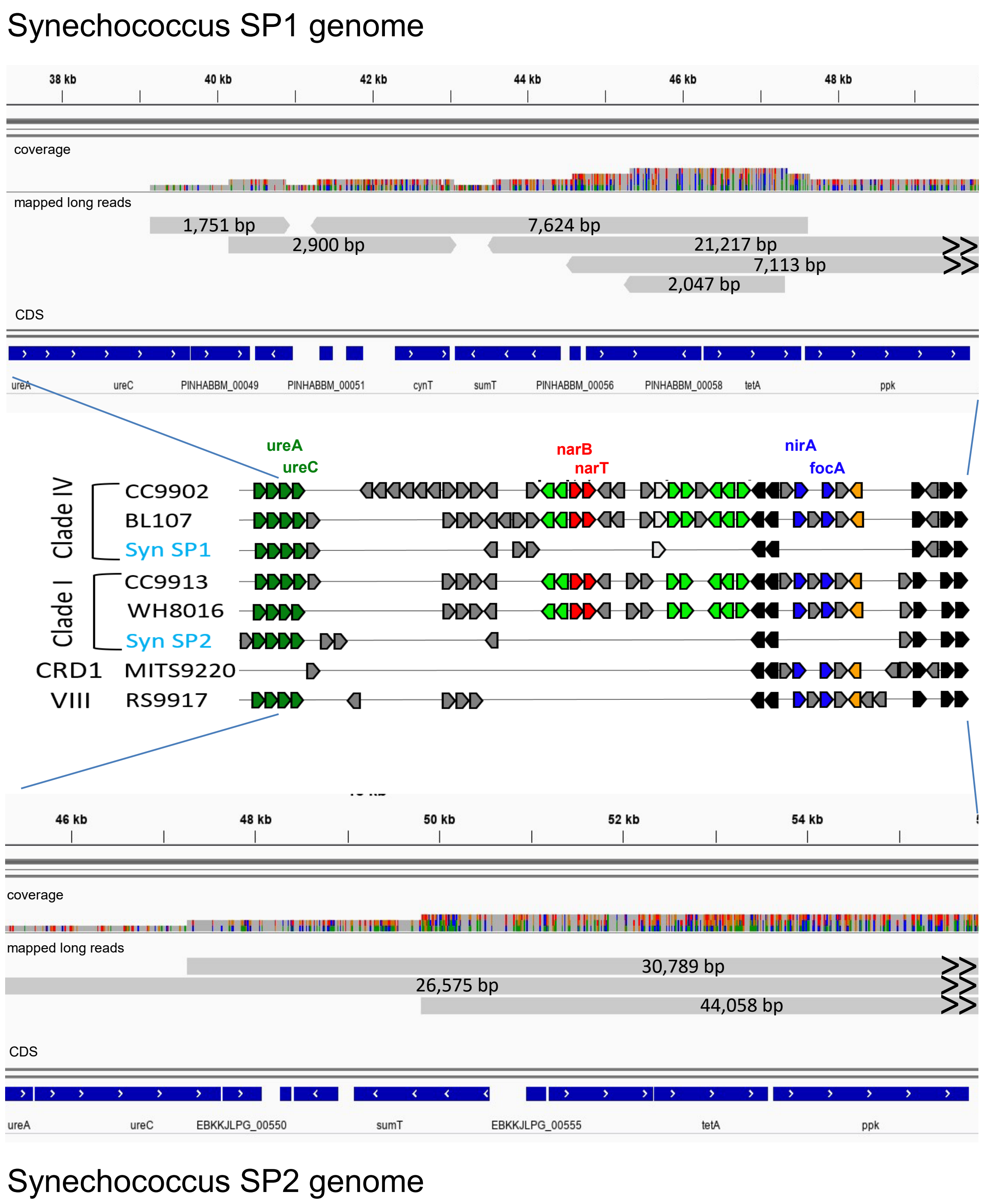


Figure S4. Nanopore long read coverage of the N gene region of the Synechococcus SP1 and SP2 MAGs. Arrows on the reads indicate that the read also maps to adjacent regions of the genome.

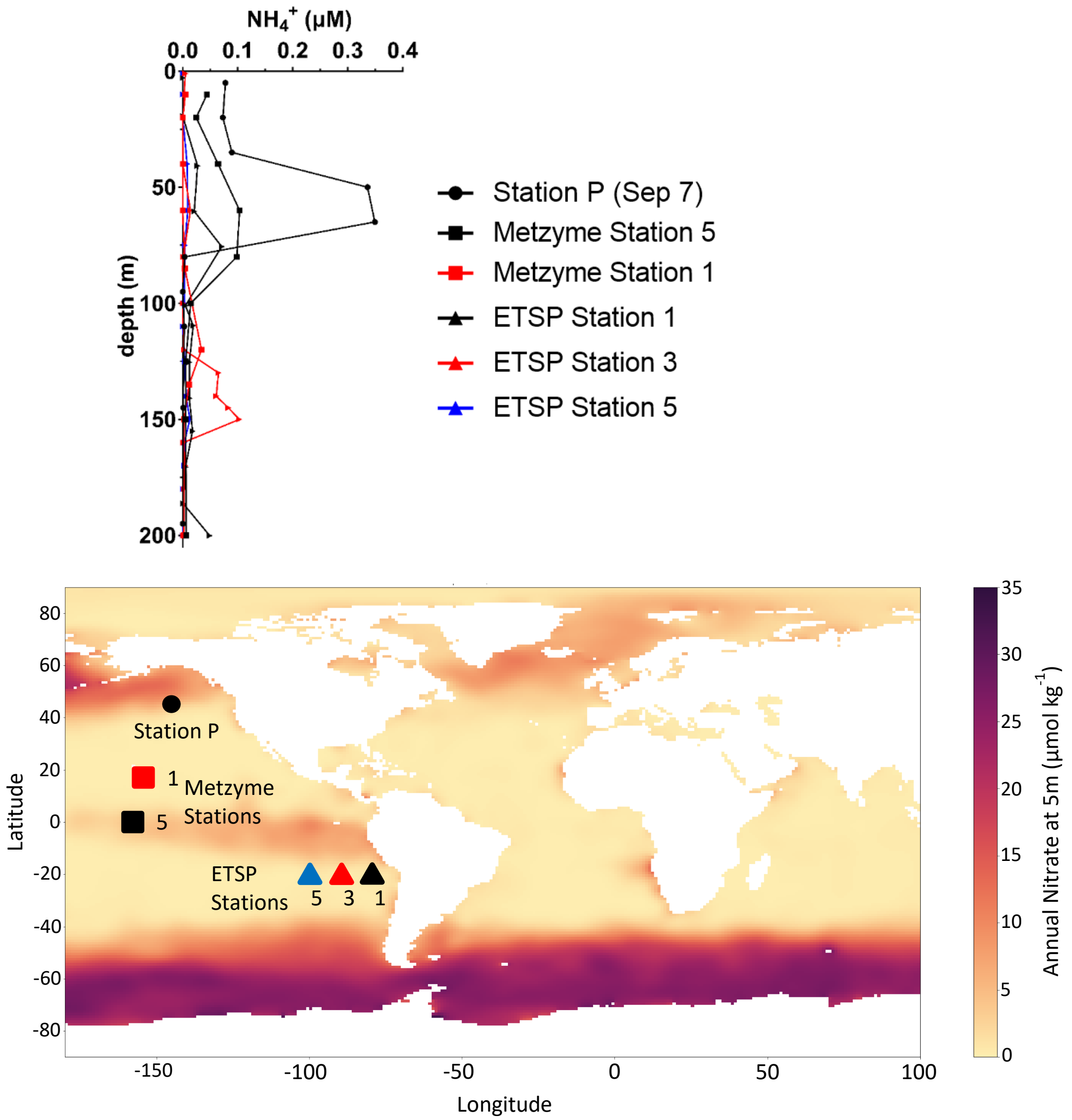


Figure S5. Comparison of ammonium concentration depth profiles between the Northeastern Pacific (Station P September 7th), the Central Equatorial Pacific, and the Southeastern Pacific. The Metzyme cruise stations are located at the equator (Station 1, 0 degrees latitude) and in the Northern Tropical Pacific (Station 5, 8 degrees latitude) (data from BCO-DMO database Project#2236 Dataset#646115) (30). Eastern Tropical South Pacific Stations (ETSP) are all located on a horizontal transect at -20 degrees latitude (-70, -80, and -100 degrees longitude for Stations 1, 3, and 5 respectively) (data from BCO-DMO database Project#555516 Dataset#820165) (31).

**Citations**


1. D. A. Siegel, et al., An operational overview of the EXport Processes in the Ocean from RemoTe Sensing (EXPORTS) Northeast Pacific field deployment. Elem Sci Anth 9, 00107 (2021).
2. J. R. Graff, M. J. Behrenfeld, Photoacclimation Responses in Subarctic Atlantic Phytoplankton Following a Natural Mixing-Restratification Event. Frontiers in Marine Science 5 (2018).
3. S. Andrews, Others, FastQC: a quality control tool for high throughput sequence data (2010).
4. A. M. Bolger, M. Lohse, B. Usadel, Trimmomatic: a flexible trimmer for Illumina sequence data. Bioinformatics 30, 2114–2120 (2014).
5. J. Zhang, K. Kobert, T. Flouri, A. Stamatakis, PEAR: a fast and accurate Illumina Paired-End reAd mergeR. Bioinformatics 30, 614–620 (2014).
6. B. Buchfink, C. Xie, D. H. Huson, Fast and sensitive protein alignment using DIAMOND. Nat. Methods 12, 59–60 (2015).
7. W. De Coster, S. D'Hert, D. T. Schultz, M. Cruts, C. Van Broeckhoven, NanoPack: visualizing and processing long-read sequencing data. Bioinformatics 34, 2666–2669 (2018).
8. S. Nurk, D. Meleshko, A. Korobeynikov, P. A. Pevzner, metaSPAdes: a new versatile metagenomic assembler. Genome Res. 27, 824–834 (2017).
9. B. Langmead, S. L. Salzberg, Fast gapped-read alignment with Bowtie 2. Nat. Methods 9, 357–359 (2012).
10. H. Li, et al., The Sequence Alignment/Map format and SAMtools. Bioinformatics 25, 2078–2079 (2009).
11. D. D. Kang, et al., MetaBAT 2: an adaptive binning algorithm for robust and efficient genome reconstruction from metagenome assemblies. PeerJ 7, e7359 (2019).
12. Y.-W. Wu, Y.-H. Tang, S. G. Tringe, B. A. Simmons, S. W. Singer, MaxBin: an automated binning method to recover individual genomes from metagenomes using an expectation-maximization algorithm. Microbiome 2, 26 (2014).
13. J. Alneberg, et al., CONCOCT: Clustering cONtigs on COverage and ComposiTion. arXiv [q-bio.GN] (2013).
14. C. M. K. Sieber, et al., Recovery of genomes from metagenomes via a dereplication, aggregation and scoring strategy. Nat Microbiol 3, 836–843 (2018).
15. D. H. Parks, M. Imelfort, C. T. Skennerton, P. Hugenholtz, G. W. Tyson, CheckM: assessing the quality of microbial genomes recovered from isolates, single cells, and metagenomes. Genome Res. 25, 1043–1055 (2015).
16. P. A. Chaumeil, A. J. Mussig, P. Hugenholtz, D. H. Parks, GTDB-Tk: a toolkit to classify genomes with the Genome Taxonomy Database. Bioinformatics (2019).
17. T. J. Treangen, D. D. Sommer, F. E. Angly, S. Koren, M. Pop, Next generation sequence assembly with AMOS. Curr. Protoc. Bioinformatics Chapter 11, Unit 11.8 (2011).
18. H. Li, Minimap2: pairwise alignment for nucleotide sequences. Bioinformatics 34, 3094–3100 (2018).
19. J. T. Robinson, et al., Integrative genomics viewer. Nat. Biotechnol. 29, 24–26 (2011).
20. A. M. Eren, et al., Anvi'o: an advanced analysis and visualization platform for 'omics data. PeerJ 3, e1319 (2015).
21. D. Hyatt, et al., Prodigal: prokaryotic gene recognition and translation initiation site identification. BMC Bioinformatics 11, 119 (2010).
22. D. H. Parks, et al., A complete domain-to-species taxonomy for Bacteria and Archaea. Nat. Biotechnol. 38, 1079–1086 (2020).
23. B. M. Satinsky, S. M. Gifford, B. C. Crump, M. A. Moran, Use of internal standards for quantitative metatranscriptome and metagenome analysis. *Methods Enzymol.* **531**, 237–250 (2013).

24. S. M. Gifford, *et al.*, Microbial Niche Diversification in the Galápagos Archipelago and Its Response to El Niño. *Front. Microbiol.* **11**, 575194 (2020).
25. A. C. E. Darling, B. Mau, F. R. Blattner, N. T. Perna, Mauve: multiple alignment of conserved genomic sequence with rearrangements. Genome Res. 14, 1394–1403 (2004).
26. C. L. M. Gilchrist, Y.-H. Chooi, Clinker & clustermap.js: Automatic generation of gene cluster comparison figures. Bioinformatics (2021) https:/doi.org/10.1093/bioinformatics/btab007.
27. O. Aumont, C. Ethé, A. Tagliabue, L. Bopp, M. Gehlen, PISCES-v2: an ocean biogeochemical model for carbon and ecosystem studies. Geosci. Model Dev. 8, 2465–2513 (2015).
28. C. Richon, A. Tagliabue, Biogeochemical feedbacks associated with the response of micronutrient recycling by zooplankton to climate change. *Glob. Chang. Biol.* **27**, 4758–4770 (2021).
29. U. Omasits, C. H. Ahrens, S. Müller, B. Wollscheid, Protter: interactive protein feature visualization and integration with experimental proteomic data. *Bioinformatics* **30**, 884–886 (2014).
30. M. A. Saito, Nutrients, targeted proteomics, and pigments from the METZYME cruise (KM1128). Biological and Chemical Oceanography Data Management Office (BCO-DMO) Version Date 2018-05-24 (2018).
31. K. L. Casciotti, A. E. Santoro, A. N. Knapp, Dissolved nitrite and ammonium concentration data from R/V Atlantis (AT15-61) cruise in Jan-Feb 2010 and R/V Melville (MV1104 cruise in Mar-Apr 2011 in the Eastern Tropical South Pacific. Biological and Chemical Oceanography Data Management Office (BCO-DMO) Version Date 2020-08-06 (2021).
